# Barcoded Rational AAV Vector Evolution enables systematic *in vivo* mapping of peptide binding motifs

**DOI:** 10.1101/335372

**Authors:** Marcus Davidsson, Gang Wang, Patrick Aldrin-Kirk, Tiago Cardoso, Sara Nolbrant, Morgan Hartnor, Malin Parmar, Tomas Björklund

**Author notes:** Present address: National Institute for Viral Disease Control and Prevention, Chinese Center for Disease Control and Prevention, 102206 Beijing, China (Gang Wang). Corresponding Author Tomas Björklund, Molecular Neuromodulation, Lund University, BMC A10 22184, Lund Sweden.

## Abstract

Engineering of Adeno-associated viral (AAV) vector capsids through directed evolution has been used to generate novel capsids with altered tropism and function_1-9_. This approach, however, involves a selection process that requires multiple generations of screenings to identify real functional capsids_2-4_. Due to the random nature of this process, it is also inherently unreproducible, and the resulting capsid variants provide little mechanistic insights into the molecular targets engaged. To overcome this, we have developed a novel method for rational capsid evolution named Barcoded Rational AAV Vector Evolution (BRAVE). The key to this method is a novel viral production approach where each virus particle displays a protein-derived peptide on the surface which is linked to a unique barcode in the packaged genome_10_. Through hidden Markov model-based clustering_11_, we were able to identify novel consensus motifs for cell-type specific retrograde transport in neurons in vivo in the brain. The BRAVE approach enables the selection of novel capsid structures using only a single-generation screening. Furthermore, it can be used to map, with high resolution, the putative binding sequences of large protein libraries.

---

Here, we have developed a novel high-throughput approach to engineering of AAV capsids that encompasses all the benefits of rational design^12-16^ while maintaining the broad screening diversity permitted by the directed evolution. This is achieved using only a single screening step. In the first application of BRAVE screening, we utilize peptides derived from proteins with documented synapse interaction with the aim to develop novel AAV capsid variants that are efficiently taken up and transported retrogradely in neurons *in vivo*. This combinatorial method enabled the generation and simultaneous functional mapping of close to 4 million unique AAV virion variants in parallel *in vitro* and *in vivo*. In a single round of screening, we selected 25 candidates, all of which could be derived into functional viruses with the capacity to become retrogradely transported *in vivo* or efficiently transduce of neurons *in vitro*. We identified one novel capsid variant expressing highly efficient retrograde transport in both rat and human dopamine (DA) neurons *in vivo,* and a panneuronal retrogradely transported AAV which we utilized to elucidate the function of basolateral amygdala projections to the dorsal striatum.

To link capsid structure to an *in vivo* expressed molecular barcode, we generated a novel AAV production plasmid. A barcode is inserted in the 3’ UTR of GFP of a gutted self-complementary AAV genome^17, 18^ and the AAV2 Rep/Cap genes are expressed from the same plasmid, creating a linkage between capsid structure and barcode. Here, peptides are displayed at N587 of VP1 capsid protein^13^ (Supplementary Fig. S1). This mutates the wild-type AAV2 heparan sulfate proteoglycan binding motif^14^. The AAV virus produced from this plasmid with no inserted peptide is hereafter referred to as MNMnull.

With the aim to targeted neurons at their terminals, we selected 131 proteins based on their documented affinity to synapses (Fig. 1A and Supplementary Fig. S2 and S3). Their amino acid (aa) sequences were computationally digested into overlapping 14aa or 22aa long polypeptides and three alternative linkers were then added to the 14 aa polypeptides (1-2. in Fig. 1B). The resulting 92 358 oligonucleotides were synthesized in parallel on an array (3. in Fig. 1B). They were then assembled into the backbone plasmid to allow for packaging of replication deficient AAV viral particles, where the peptide is displayed on the capsid surface and a 20bp random molecular barcode (BC) is included as part of the genome (4. in Fig. 1B and Supplementary Fig. S1). In parallel, the same plasmid library was utilized to generate a Look-Up table (LUT), linking the random barcodes to the respective peptide^10^ (5a. in Fig. 1B and Supplementary Fig. S4 and S14). The resulting library contained 3 934 570 unique combinations of peptide and barcode, with 50-fold oversampling of the peptides (essential for noise filtration, and mapping of oligonucleotide array-induced mutations).

**Figure 1.**
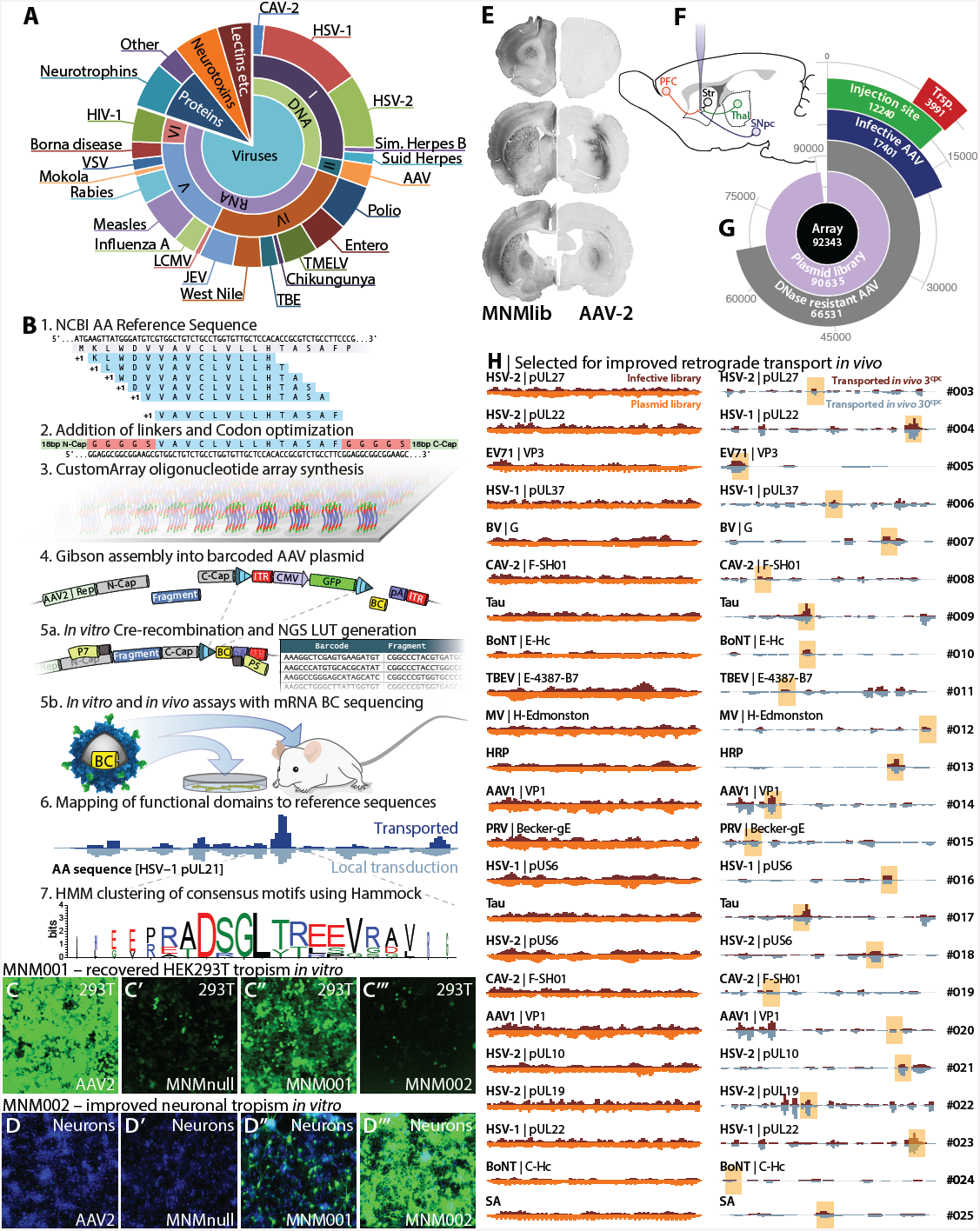
Assessment of retrograde transport in the brain using the BRAVE AAV capsid library approach. **(A)**Pie-chart displaying the selected 131 proteins with documented affinity to synapses.**(B)**Schematics of BRAVE procedure; **1.** NCBI reference amino acid (aa) sequences were computationally digested into 14aa (or 22aa) long polypeptides with a 1aa shifting sliding window. **2.** Three alternative linkers were added to the 14aa polypeptides e.g., a rigid linker with 5 Alanine residues (14aaA5). **3.** 92,358 codon optimized sequences were synthesized in parallel on an oligonucleotide array. **4.** The pool of oligonucleotides was assembled into a novel AAV production backbone with *Cis-*acting AAV2 Rep/Cap and ITR-flanking CMV-GFP. A 20bp random molecular barcode (BC) was simultaneously inserted in the 3’ UTR of the GFP gene. **5a.** Using a fraction of the plasmid prep, exposed to Cre-recombinase *in vitro*, a look-up table (LUT) was generated linking BC to peptide. **5b.** Using the same plasmid pool from step 4, an AAV was produced (MNMlib) and multiple parallel screening experiments were performed both *in vitro* and *in vivo* followed by BCs sequencing from mRNA **6.** Through the combination of the sequenced barcodes and the LUT, efficacy can be mapped back to the original 131 proteins and consensus motifs can be determined using the Hammock, hidden Markov-model based clustering approach (**7.**). (**C-C’’’**) First proof-of-concept study using BRAVE to re-introduce tropism for HEK293T cells *in vitro*. Wild-type AAV2 displays very high infectivity attributed to Heparin Sulfate (HS) proteoglycan binding (**C**). The MNMnull serotype disrupts this binding (**C’**). The first novel capsid generated from single-generation BRAVE screening in HEK293T, MNM001, displayed a significantly recovered tropism (**C’’**). (**D-D’’’**) In a second experiment, we used the BRAVE technology to improve the infectivity of primary cortical rat neurons *in vitro*. Both AAV2-WT and the MNMnull vector displays very poor infectivity of primary neurons (**D-D’**) and the MNM001 displays some improvement (**D’’**). BRAVE screening in primary neurons identified a number of peptides clustering over a C-terminal region of the HSV-2 pUL1 protein which improved the infectivity of primary neurons in culture dramatically (**D’’’**). (**E**) *In vivo* expression pattern of GFP after injection of 30cpc library into the rat forebrain compared to an AAV2-WT vector at the same titer.(**F**) Connectivity diagram showing the injection site (striatum, Str) and connected neuronal populations frontal cortex (PFC), thalamus (Thal) and substantia nigra (SNpc) utilized for BRAVE screening *in vivo* for retrograde transport. (**G**) Polar plot showing the absolute quantities of the unique peptides recovered at each step of the BRAVE assay. (**H**) From the *in vivo* screening for improved retrograde transport capacity, we selected 23 peptides from 20 proteins that all were represented by multiple barcodes and found in multiple animals (See Supplementary Document S14). 23 of the 25 *de novo* AAV capsid structures allowed for higher than or at par with AAV2-WT packaging efficacy (Supplementary Fig. S1), all with retrograde transport ability (Supplementary Fig. S7).Davidsson *et al.,* 2018. *In vivo* single-generation AAV-screen

To ensure that each virion is assembled using only one mutated capsid protein variant and that the corresponding barcode is packaged inside, the AAV library plasmid was supplied at very low concentrations during production^6, 19^, 3 or 30 copies per cell (cpc, Supplementary Fig. S4). Using the AAV vector library, we performed multiple parallel screening experiments *in vitro* and *in vivo*, followed by sequencing of barcodes expressed in the mRNA. Efficacy could be mapped back to the original 131 proteins with the help of the LUT and consensus motifs determined using the Hammock method (6-7. in Fig. 1B). We utilized the BRAVE technology to screen for the re-introduction of tropism for HEK293T cells *in vitro* (Fig. 1C) which was lost when the HS binding motif was removed in the MNMnull capsid (Fig. 1C-C’). In the screening of the 4 million uniquely barcoded capsid variants, we found several regions from the 131 included proteins that conferred a significantly improved infectivity over the parent MNMnull capsid structure (Supplementary Fig. S5, S6 and S14). One peptide from HSV-2 surface protein pUL44 was selected and a first novel capsid variant was generated (MNM001) (Supplementary Fig. S6). This indeed displayed a recovered tropism to the HEK293T cells (Fig. 1C’’). Through a second BRAVE screen in primary cortical neurons, we identified a number of peptides clustering over a C-terminal region of the HSV-2 pUL1 protein (Supplementary Fig. S6). From these data, we generated a novel AAV capsid (MNM002) which improved the infectivity of primary neurons in culture dramatically compared to both the AAV2-WT and the MNMnull vector (Fig. 1D-D’’’ and Supplementary Fig. S6 and S14).

To identify individual novel AAV capsid variants with efficient retrograde transport in neurons *in vivo*, the AAV libraries (MNMlib[Xcpc]) were injected into the forebrain of adult rats. Compared to the standard AAV2-WT vector, the inserted peptides confer a striking change in the transduction pattern with both broader spread of transduction and retrograde transport to the connecting afferent regions (Fig. 1E). Eight weeks after injection, total RNA was extracted from the injection site and from three connected regions (Fig. 1F) and the transcribed barcodes sequenced (Supplementary Document S14). Analysis of the unique peptides (identified using the barcodes and the LUT) throughout the BRAVE pipeline provides unique insights into the efficacy of the process (Fig. 1G). More than 98% of the designed oligonucleotides (Array, black circle) was successfully cloned into the plasmid library and barcoded (purple ring) and 72% of the peptides permitted complete assembly of the AAV (as assessed by DNase treatment, gray ring in 1G). Barcodes recovered at the dissected regions reveal that ≈ 13% of the inserted peptides promoted efficient transduction in the brain (green ring segment) and ≈ 4% promoted retrograde transport in neurons (red ring segment in 1G). Pooling all experiments including *in vitro* experiments revealed that at least 19% of the peptides could promote infectivity in cell types tested (dark blue ring segment in 1G).

From this large *in vivo* BRAVE screening, we selected 23 peptides from 20 proteins that all were represented by multiple barcodes and found in multiple animals (Fig. 1H and Supplementary Fig. S5, S7 and S14). Twenty-one of the 23 *de novo* AAV capsid structures found *in vivo* (and the two from the *in vitro* screening) allowed for higher than or on par with AAV2-WT packaging efficacy (Supplementary Fig. S1). Using a hidden Markov model (HMM) based clustering^11^, of all peptides with a retrograde transport ability, we could determine the putative consensus motif for each of the 23 serotypes, providing a foundation for directed optimization (Supplementary Document S14).

In the BRAVE design, we can compare the function of peptides across multiple regions of the brain and also between animals (Supplementary Document S14). This allows us to map domains of proteins that promote specific transport or uptake abilities. Exemplified by HSV pUL22 protein, peptides throughout the protein can drive uptake at the injection site in the striatum (Fig. 2A). However, only a C-terminal region of the protein displayed reproducible transport to all afferent regions; cortex, thalamus and substantia nigra. HMM clustering of all peptides recovered at these sites revealed two overlapping consensus motifs (Fig. 2B-D). Those were generated into the MNM004 and MNM023 (Fig. 2C) capsid structures, respectively, with both displaying similar transport patterns *in vivo* (Supplementary Fig. S7E versus S7X). The MNM004 capsid promoted efficient retrograde transport to all afferent regions as far back as the medial entorhinal cortex (Fig. 2E and Supplementary movie 1). The parent AAV2-WT capsid promoted efficient transduction at the site of injection but very little retrograde transport of the vector (Fig. 2E). While this manuscript was finalized, another peptide, LADQDYTKTA, was published promoting strong retrograde transport when displayed in the same location on the AAV2 capsid surface (AAV2-Retro)^9^. When compared *in vivo*, the two vectors displayed very similar retrograde transport properties with the MNM004 displaying equal or higher transport efficacy (Fig. 2F-L and Supplementary Fig. S8).

**Figure 2.**
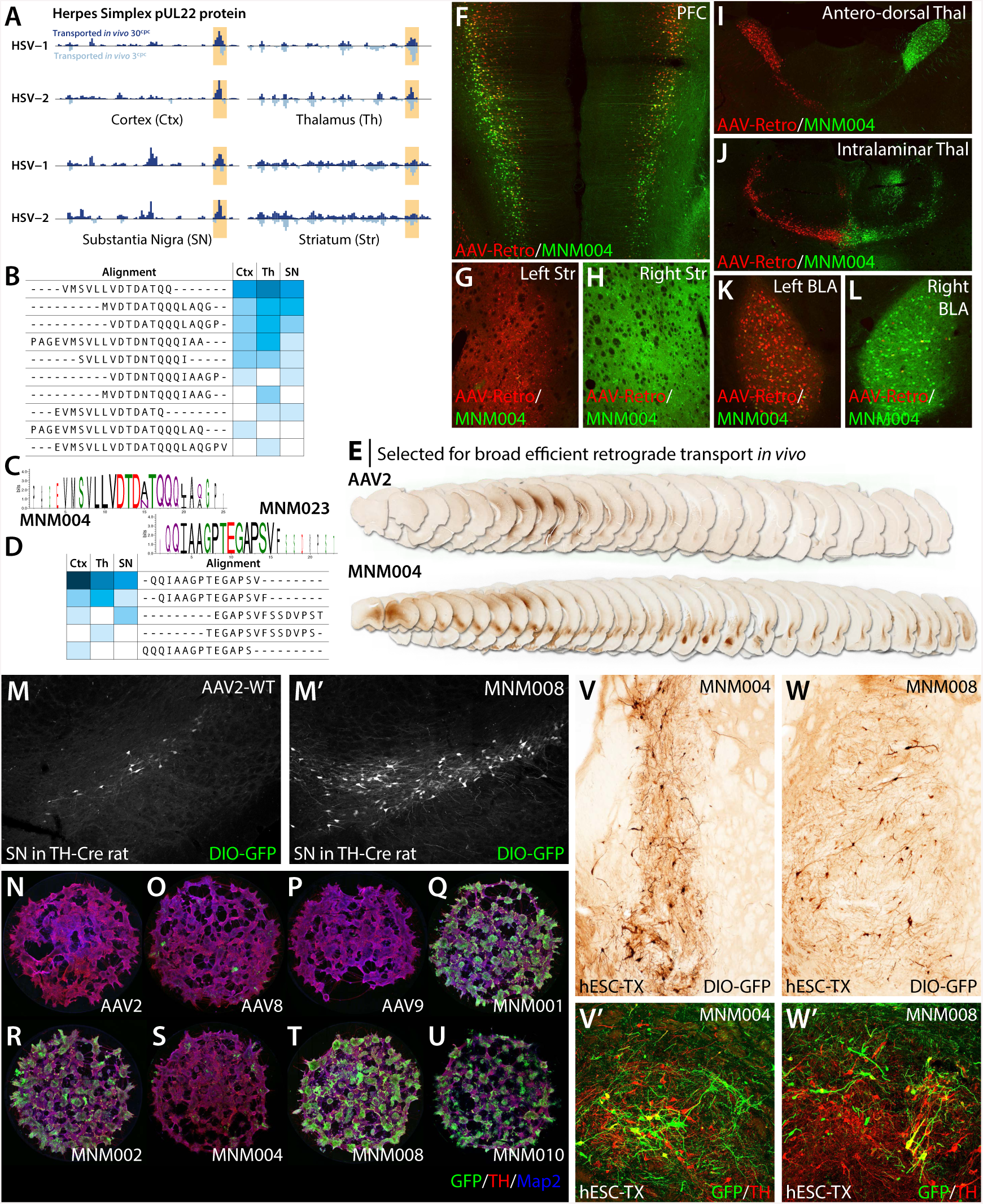
Characterization of the MNM004 capsid for retrograde transport *in vivo* and generation of capsids infecting DA neurons. (**A**) A C-terminal region of the HSV pUL22 protein (highlighted), displayed reproducible transport to all afferent regions while not showing the same bias at the injection site (striatum). (**B-D**) HMM clustering of all peptides displaying these properties revealed two overlapping consensus motifs (**C**). Those were generated into the MNM004 (**B**) and MNM023 (**D**) AAV capsid variants respectively. (**E**) *In vivo* comparison between MNM004 and the parent AAV2 after unilateral striatal injection. Both vectors express GFP and sections were developed using brown DAB-peroxidase reaction (See Supplementary movie 1 for a 3D visualization). (**F-L**) Same animal comparison between MNM004 and AAV2 capsid with the previously published LADQDYTKTA peptide (AAV2-Retro). AAV2-Retro express mCherry and was injected into the left striatum (**G**) while MNM004 express GFP and was injected into right striatum (**H**) at matched titers. Monitored afferent regions include pre-frontal cortex (PFC; **F**), antero-dorsal (**I**) and intralaminar thalamic nuclei (**J**) and the basolateral amygdala (BLA; **K-L**) (see Supplementary Fig. S8 for additional information). (**M-M’**) *In vivo* comparison between the novel CAV2-derived MNM008 and AAV2-WT assessing infectivity of dopamine neurons from their terminals in the striatum. Both vectors express Cre-inducible GFP (DIO-GFP) and were injected into the striatum of TH-Cre knock-in rats. (**N-U**) Assessment of retained neuronal tropism in hESC-derived DA neuroblasts *in vitro*. (**V-W’**) Assessment of retrograde infectivity in humanized rats. These animals first received a human embryonic stem cell (hESC) derived DA-rich neuronal transplant (expressing Cre) into the striatum. Six months later, the MNM008 or MNM004 vectors (expressing DIO-GFP), were injected into the frontal cortex.

In a final BRAVE screening experiment, we aimed to develop a novel AAV capsid variant with efficient retrograde transport to dopamine neurons of the substantia nigra from injections into the striatal output region. In this screening, we identified two regions of the CAV-2 capsid protein (often-used viral vector for the targeting of DA neurons from the terminal *in vivo*^20)^ in close proximity (Supplementary Document S14). One motif shared homology with the lectin soybean agglutinin (SA, also used as a retrograde tracer^21^). Using a Cre-inducible AAV genome (CMV-loxP-GFP) injected into the striatum of TH-Cre knock-in rats, we found that the CAV-2-based MNM008 improved the retrograde infectivity of nigral neurons dramatically from single neurons in the SN in AAV2-WT to the majority (Fig. 2M-M’). A major drawback of *de novo* capsid design using animal models has been the lack of predictability with regards to the translation to human cells. The BRAVE approach is expected to improve this through the use of naturally occurring peptides from viruses and proteins with known function in the human brain. To establish that the properties of MNM008 was not limited to rodent DA neurons, we then performed an experiment on animals transplanted with intranigral transplants of Cre-expressing human embryonic stem cell (hESC) derived-DA neurons ^22^. Six months after the transplantation, the MNM008 vector was injected into the frontal cortex of the animals. This was efficiently transported back to the transplanted neurons innervating this region (Fig. 2V-V’), with the majority of these cells being DAergic (Fig. 2V’). Interestingly, the MNM004 which does not have this ability in the rat brain showed similar potency in the transplanted human neurons (Fig. 2W-W’). Using the same *in vitro* hESC differentiation protocol we confirmed that the *de novo* capsid variants (MNM002, 008 and 010) which displayed high tropism on primary rodent neurons also showed much higher tropism than the wild-type AAV-variants (Fig. 2N-U’ and Supplementary Fig. S11) in human neurons *in vitro*. Of note is that the MNM004 capsid variant, so efficient *in vivo* was not at all suitable for *in vitro* transduction (Fig. 2S).

The BRAVE approach provides a unique possibility to systematically map protein function. We therefore utilized this approach to display peptides from endogenous proteins involved in Alzheimer’s disease; APP and microtubule associated protein Tau to provide insights into the mechanism underlying the proposed cell-to-cell communication of these proteins in the disease^23, 24^. In the mapping of APP we found two regions that conferred retrograde transport, one in the sAPP N-terminal region and one in the Amyloid beta region (Supplementary Fig. S9). The functional properties of peptides originating from Tau were even more striking. In this protein, a central region conveyed very efficient retrograde transport (Fig. 3A & Supplementary Fig. S9, S14). Two novel capsid structures were generated from this region, MNM009 and MNM017. Both capsids promoted retrograde transport *in vivo* (Supplementary Fig. S7J and R) but MNM017 also displayed additional interesting properties. MNM017 infected both primary neurons and primary glial cells *in vitro* with very high efficacy (Fig. 3B, Supplementary Fig. S6 and S10), including human primary glia (Fig. 3C-C’’). Using this property, we then performed a displacement experiment comparing the MNM017 to the neurotrophic MNM002 capsid which is not generated from a Tau-derived peptide (Fig. 3D and Supplementary Fig. S10). Three groups of primary neuron populations were pre-treated with different recombinant Tau variants (T44, T39 and K18). The T44 variant had no apparent effect on the MNM017 to MNM002 ratio of infectivity while K18 enhanced the infectivity of the MNM017 and the T39 variant efficiently blocked the infectivity compared to MNM002 (Fig. 3D). This suggests that the Tau peptide is utilizing a receptor on the neurons that also has a binding activity of full length human Tau protein.

**Figure 3.**
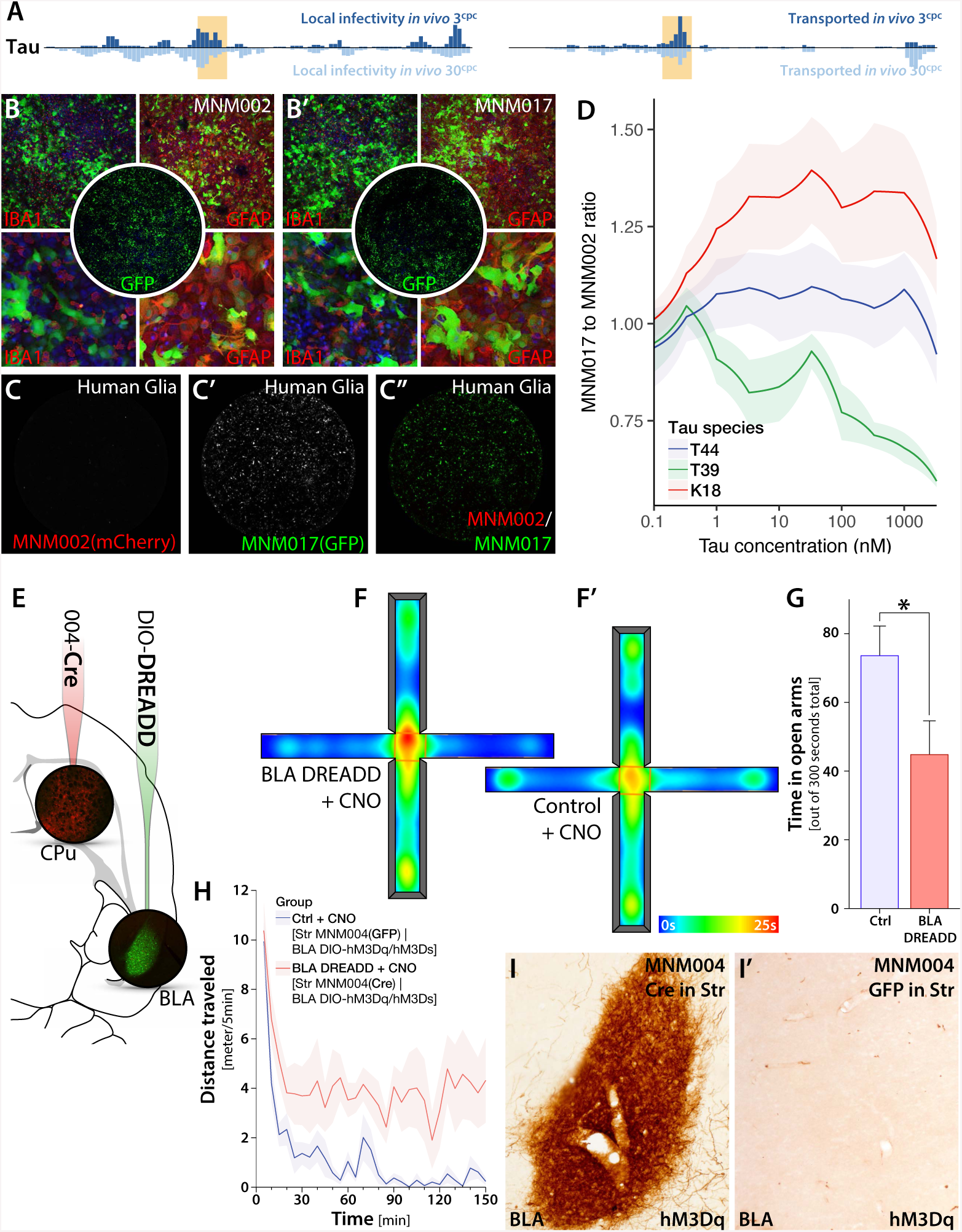
Mapping of proteins involved in Alzheimer’s disease and functional dissection of the basolateral amygdala. (**A**) Peptides originating from a central region (highlighted) in the microtubule associated protein Tau were found to convey both a very efficient retrograde transport and local infectivity in the rat brain. Two novel capsid variants were generated from this region, MNM009 and MNM017. (**B-B’**) *In vitro* assessment of two *de novo* capsid variants (MNM002 & MNM017) in primary glial cells. **(C-C’’)** *In vitro* transduction of human primary Glia using MNM002 and MNM017. (**D**) Recombinant Tau displacement experiment *in vitro* of MNM017 compared to the MNM002 capsid which is not generated from a Tau-related peptide. Primary cortical neurons were first treated with different recombinant Tau species at varying concentration and then infected by MNM017(GFP) and MNM002(mCherry) simultaneously. (**E**) Experimental injection paradigm for functional assessment of the afferents from the basolateral amygdala (BLA) to the dorsal striatum (Cpu). We injected the Cre-expressing MNM004 vector in the Cpu and a Cre-inducible (DIO) chemogenetic (DREADD) vector into the BLA bilaterally into wild-type rats. (**F-G**) Assessment of fear and anxiety phenotype using the elevated plus maze (EPM) where the animals spent significantly less time on the open arms (**F**) compared to control animals (**F’)**, where the Cre gene was replaced with GFP. (**H**) The increased anxiety phenotype was accompanied by significant hypermobility in the open-field arena and a fear phenotype including excessive digging, severe sweating and episodes of freezing (See Supplementary movie 2). (**I-I’**) Visualization of selective DREADD expression in the BLA using immunohistochemistry together with brown DAB-peroxidase precipitation reaction.

The BRAVE generated capsid variants provide a powerful tool for brain connectivity studies. We utilized the MNM004 capsid variant to answer an outstanding question regarding the functional contribution of the afferents from the basolateral amygdala (BLA) to the dorsal striatum. This was conducted using a retrogradely-induced chemogenetics (DREADD) approach (Fig. 3E). After selective induction of activity of the BLA neurons projecting to the dorsal striatum, we found a striking fear and anxiety phenotype (Fig. 3 F-G) compared to control animals (where the Cre gene was replaced with GFP). This stands in stark contrast to the function of the BLA projections to the ventral striatum shown to promote positive stimuli^25^. This increased anxiety phenotype was accompanied by significant hypermobility (Fig. 3H and Supplementary Fig. S12) and a fear phenotype including excessive digging, severe sweating and episodes of freezing (See Supplementary movie 2). *Post mortem* immunohistochemistry confirmed a very selective DREADD expression in the BLA (Fig. 3I-I’).

While recent additions of Cre-recombinase restricted selection and deep sequencing for optimization has improved the accuracy and potency of directed evolution^3^, it is still hampered by the difficulty in extrapolating the findings to other strains or species than the one used for screening^26, 27^. An alternative approach is rational design where systematic changes are made based on the known properties of the capsid, through systematic amino-acid substitutions or display of high affinity nanobodies of the capsid surface^12-16^. Although functionally more stringent, this approach provides less diversity and has more restricted functional potential.

The BRAVE screening approach for technology is unique in that it allows for a single-generation screening and systematic mapping of capsid motifs. It also provides several key improvements; the use of barcode in the packaged genome allows for removal of Cap gene which ensures that serial infectivity cannot occur during the production. The expression of the barcode in mRNA allows for detection of single virions transported to the target region without relying on Adenoviruses for *in vivo* recovery.

The use of a stable bacterial library together with oversampling of barcodes allows for a very efficient error control *in vivo* based solely on unique barcode count (not mRNA expression level) and number of animals displaying the same function for one exact peptide. This provides unprecedented accuracy to the screening and removes the need for multiple generations of screening. Going forward, the LUT and *in vivo* data can be re-used to further improve on target selection and filter out off-target infectivity. Furthermore, the single-generation BRAVE screening has the potential to be used together with both single-cell and *in situ* sequencing approaches^28^. This opens up for the development of highly selective capsid structures with a large potential for clinical applications and brain connectivity studies.

## Availability of data and materials

The datasets supporting the conclusions of this article will available in the NCBI Sequence Read Archive (SRA) with the Accession numbers: SRP149133 The R-based workflow is publicly available as a Git repository at https://bitbucket.org/MNM-LU/aav-library and as a Docker image: Bjorklund/aavlib:v0.2.

## Authors’ contributions

TB, MD, GW and PA designed the experiment; MD, GW, PA, TC, SN, and MH performed the wet experiments; TB analyzed the sequencing data; MD, PA, GW, MP and TB wrote the manuscript.

## Acknowledgement

The authors would like to thank the staff at the National Genomics Infrastructure (NGI) of SciLifeLab, Sweden and UCLA Clinical Microarray Core, USA for expert assistance in the sequencing performed using the Illumina NextSeq technology. The authors would also like to thank Anna Hammarberg for assistance and much appreciated help with *in vitro* fluorescence imaging. pscAAV-GFP was a gift from John T Gray (Addgene plasmid # 32396), pHGTI-adeno was a gift from Julie Tordo and recombinant Tau was a gift from Virginia Lee and Alexander Crowe.

## Document S14 | Bioinformatics output

This document contains the complete formatted output of the bioinformatics pipeline. The datasets required to re-run this analysis pipeline are available in the NCBI Sequence Read Archive (SRA) with the Accession numbers: SRP149133 The R-based workflow is publicly available as a Git repository at https://bitbucket.org/MNM-LU/aav-library and as a Docker image: Bjorklund/aavlib:v0.2.

## Online materials and methods

### AAV library backbone plasmid cloning

The backbone plasmid used for cloning the barcoded modified AAV capsids was developed from a self-complementary AAV (scAAV) vector expressing GFP (pscAAV-GFP^29^) together with components from pDG^30^ (Adenovirus genes VA, E2A and E4 are deleted). The final plasmid contained an eGFP expression cassette driven by a CMV promoter and the wild-type AAV2 Rep/Cap genes controlled by the mouse mammary tumor virus (MMTV) promoter. First, the XbaI, BsiWI and MluI sites were inserted between XhoI and HindIII sites in pscAAV-GFP [Addgene #32396], by annealing Primer 1 and Primer 2 (P1&P2) (for primers see Supplementary Fig. S13) and ligating it into pscAAV-GFP. Second, an NheI site was introduced between sequences of N587 and R588 of VP1 capsid protein by overlap extension PCR^31^ using modified pDG as template and primers P3-P6. Primers P3-P4 and P5-P6 were used in the first round of PCR and primers P3-P6 were used in the second round of PCR. Beside the NheI site, the final PCR product also contained a BsiWI and a MluI site to facilitate subsequent cloning and LoxP-JTZ17 insertion (for Cre recombination and next-generation sequencing of the final library). Finally, the modified pscAAV-GFP was digested using XbaI and MluI, the overlap extension PCR product was digested with MluI and BsiWI, and pDG was digested by BsiWI and XbaI. The three DNA fragments were then ligated to acquire the final backbone plasmid for the AAV library.

### Selecting proteins for peptide display

Candidates of peptides to be inserted were derived from known neuron-related proteins. 131 proteins were selected, belonging to five categories; neurotropic viruses, lectins, neurotrophins, neurotoxins, and neuronal proteins. The candidate protein selection was based on known interaction between the proteins and neurons in binding and different stages of AAV infection and replication process (e.g., internalization, endosomal trafficking, nuclear import, etc.). See Supplementary Fig. S3 for the complete list. Peptides were designed to be incorporated between N587 and R588 of VP1 capsid protein^13^, a site that previously was reported to tolerate insertion of large peptides^16, 32^ and blocks heparan sulfate proteoglycan binding^14, 15^. Four different peptide conformations were designed as: A-14aa-A, A-22aa-A, A5-14aa-A5, G4S-14aa-G4S (For oligos see Supplementary Fig. S13). They contained two lengths of peptides of 14 or 22 amino acid (aa) residues and were flanked by either a spacer of one amino acid of alanine (A) or a short linker (A5 or G4S)^33,34^. All possible unique peptides of 14 or 22aa from the candidate proteins were identified and generated by a sliding window approach using a purpose-built R workflow (see Availability of data and materials section and Supplementary Document S14). The peptide library was reverse translated to oligonucleotides using human codon optimization for high-level expression in HEK293 cells.

### Array oligonucleotide synthesis and amplification

The final oligonucleotide pool containing 92,918 unique oligonucleotides was synthesized using 90k DNA array (Custom Array), which encoded all possible unique peptides from A-14aa-A, and every third selected peptides from the other three peptide conformations. The oligonucleotide pool was amplified and prepared for Gibson assembly in an emulsion PCR with long extension time to reduce PCR artifacts^10, 35^ using 32ng of oligonucleotides (1pmol) and primers P7 and P8 (for primers see Supplementary Fig. S13).

PCR mix was prepared using Phusion Hot Start II High-Fidelity DNA polymerase (Thermofisher) according to manufacturer’s recommendation with the exception of adding 0.5µg/µl BSA (NEB). Briefly, 9 volumes of an oil-surfactant mixture (92.95% of Mineral oil, 7% of ABIL WE and 0.05% of Triton X-100) was added to the PCR mixture and an emulsion was created by homogenizing for 5min at a speed of 4m/s using MP FastPrep-24 Tissue and Cell Homogenizer (MP Biomedicals). The PCR program used for the emulsion PCR was; 1 cycle of 30s at 98°C, 30 cycles of 5s at 98°C, 30s at 65°C and 2 min (8-fold of regular extension time) at 72°C and finally 1 cycle of 5min at 72°C. After the PCR, the emulsion was broken by adding 2 volumes of isobutanol to each tube. The aqueous phase containing the PCR product was separated by a short centrifugation (16,000g for 2min), 200µl TE-buffer was added to the sample and once again centrifuged (16,000g for 2min). The PCR product was finally purified using DNA Clean and Concentrator (Zymo Research).

### AAV library cloning and barcoding

Gibson assembly was used to insert the oligonucleotide pool (into the capsid gene located outside of the ITR’s) and barcodes (downstream of GFP located inside of the ITR’s) to generate a barcoded AAV plasmid library (4. In Fig. 1B and Supplementary Fig. S1). A one-cycle PCR was performed to generate barcoded fragments with overhangs for Gibson assembly by using primers P9 & P10 (for primers see Supplementary Fig. S13). Barcoded primers were ordered from Eurofins genomics, where the barcode length was 20 nucleotides and defined as ambiguity nucleotides by using the sequence V-H-D-B (IUPAC ambiguity code) repeated five times and flanked by static sequences for primer binding. Oligos also contained a LoxP-JTZ17 site, for facilitating subsequent Cre recombination. The AAV library backbone plasmid was digested with NheI and BsrGI to generate two fragments, which had sequences compatible for Gibson assembly with the oligonucleotide pool and barcoded PCR fragments. A 40µl Gibson Assembly reaction (NEB) was performed to insert oligonucleotide pool and barcoded fragments into 200ng of the vector, using a molar ratio of 1.3:1.3:1. The reaction was incubated for 1h at 50°C and purified using DNA Clean & Concentrator-5 (Zymo Research). 1µl (37.4ng) purified Gibson assembly product was transformed into 20µl MegaX DH10B T1R Electrocomp (Thermo Fischer Scientific) cells according to the manufacturer’s protocol. Five individual transformations were performed and pooled into one tube. A small fraction of the transformed bacteria was plated on agar plates to validate the transformation efficacy. 10 clones were picked from the plates and oligonucleotide and barcode insertion was validated by restriction enzyme digestion using Bsp120I, BsrGI, and SpeI (Fastdigest, Thermo Fischer Scientific). The none plated transformed bacteria were grown o.n as a maxi prep and purified using ZymoPURE Plasmid Maxiprep Kit (ZYMO Research).

### Production of AAV vector library

HEK293T cells were seeded in 175cm^2^ cell culture flasks to achieve 60-80% confluency before transfection. 25µg (3000cpc), 250ng (30cpc) or 25ng (3cpc) of the AAV plasmid library, and 46µg of pHGTI-adeno1^36^ were transfected using calcium phosphate. The molar ratio of AAV plasmid library: pHGTI-adeno1 were 1:1, 0.01:1 (30cpc), or 0.001:1 (3cpc) respectively. The 1:1 ratio was expected to receive a chimeric AAV library, in which each single particle likely contained chimeric mutation capsid proteins. The ratio of 0.01:1 and 0.001:1 was assumed to make cells receive approximately one member from the AAV plasmid library^6^ and subsequently to receive a clean AAV library, in which each single particle was consisted of same mutation capsid proteins and a consistent barcode. Viral libraries were harvested and purified using iodixanol gradient as previously described^37^. The AAV genomic titer was determined by quantification of vector DNA as described using real time PCR^38^.

### Production of selected AAV variants

HEK293T cells were seeded in 175cm^2^ cell culture flasks to achieve 60-80% confluency before transfection. Two hours before transfection, the medium was replaced with 27ml fresh Dulbecco’s modified Eagle medium (DMEM) + 10% FBS + P/S. AAV was produced using standard PEI transfection^39^ using a three-plasmid system; transfer vector, modified AAV-capsid, and pHGT-1 adenoviral helper plasmid in a 1.2:1:1 ratio. PEI and plasmids were mixed in 3ml DMEM, incubated for 15 min and then added to the cells. 16h post transfection 27ml of medium was removed and equal volume of OptiPRO serum free medium (Thermo Fischer Scientific) + P/S was added. AAVs were harvested 72h post transfection using polyethylene glycol 8000 (PEG8000) precipitation and chloroform extraction followed by PBS exchange in Amicon Ultra-0.5 Centrifugal filters (Merck Millipore)^40^. Purified AAV’s were titered using qPCR with primers specific for promoter or transgene (for primers see Supplementary Fig. S13).

### *In vitro* transduction

HEK293T cells were cultured in DMEM +10% FBS and P/S. Primary cortical neurons or primary glial cells were isolated from embryonic rat (E18) or neonatal mouse of 1 day old as previously described^41^ and cultured in Neurobasal/B27 or DMEM high glucose medium in black 96-well flat bottom culture plates (Greiner Bio One). Cells were transduced with 2×10^2^, 2×10^3^ and 2×10^4^gc/cell. AAV was added to the medium and cells were incubated overnight. AAV containing medium was replaced with fresh medium the following day. Cells were analyzed 72h post transduction using Cellomics (Thermo Fischer Scientific) and Trophos Plate Runner (Trophos).

### Research animals

Adult female Sprague Dawley rats (225-250g) were purchased from Charles River (Germany) and were housed with free access to food and water under a 12h light/12h dark cycle in a temperature-controlled room. Adult (<180 g) female athymic “nude” rats were purchased from Harlan Laboratories (Hsd:RH-*Foxn*1^rnu^) and used as recipients for transplantation. Adult TH-Cre heterozygote Sprague-Dawley rats were supplied by SAGE labs, now Horizon Discovery, TGRA8400^42^. All experimental procedures performed in this study were approved by the Ethical Committee for Use of Laboratory Animals in the Lund-Malmö region.

### Stereotaxic AAV injection

All surgical procedures were performed under general anesthesia with a 20:1 mixture of fentanyl citrate (Fentanyl) and medetomidine hydrochloride (Dormitor), injected intraperitoneally. Targeting coordinates for all stereotactic infusions were identified relative to the bregma. A small hole was drilled through the skull and the vector solutions were injected with a 25µl Hamilton syringe fitted with a glass capillary (60–80µm i.d. and 120–160µm o.d.) and connected to an automatic infusion pump. Injection was carried out either unilaterally on the right side or bilaterally. The AAV library was injected unilaterally in striatum at the following coordinates: antero-posterior (AP), +1.2mm; medio-lateral (ML), -2.4mm; dorso-ventral (DV), -5.0/-4.0mm; tooth bar, -3.2mm. Candidate AAV vectors were infused at AP, +1.2mm; ML, -2.4mm; DV, -5.0/-4.0mm and AP, + 0.0mm; ML, -3.5mm; DV, -5.0/-4.0mm; tooth bar, -3.2mm. Comparative vector analysis between MNM004 GFP and AAV2 Retro-mCherry as well as lateralized comparison between MNM004 GFP and MNM004 mCherry were injected bilaterally at AP, +1.2mm; ML, - 2.4/+2.4mm; DV, -5.0/-4.0mm. For comparative analysis of retrograde transport to midbrain dopaminergic neurons from the striatum between MNM008, AAV-Retro and AAV2, TH-Cre animals were unilaterally injected with CTE-GFP vectors at two sites at the following coordinates: at AP, +0.8/-0.2mm; ML, -3.0/-3.7mm; DV, -5.0/-5.0/-4.0mm. The BLA modulation animals were injected with the MNM004 vector at AP, +1.2mm; ML, -2.4mm; DV, -5.0/-4.0mm; tooth bar, -3.2mm and AAV-8 vectors at AP, -2.2 mm; ML, +/- 4.8mm; DV, -7.4mm; tooth bar, -3.2mm. For the AAV-library animals, each rat received 5µl of vector solutions at dose of 2.5×10^10^ or 4.4×10^8^ vector genomes of AAV library with capsid concentration of 30cpc or 3cpc for AAV plasmid library and normal amounts pHGTI-adeno1 plasmid. The candidate vector animals received 2µl of vector per infusions site while the animals in the BLA modulation animals were infused with 3µl of MNM004 in the striatum and 3µl of Cre-inducible DREADDs AAV8 DIO-hM3Dq/rM3Ds in the BLA. Comparative analysis between vectors injected in the striatum were injected with a viral dose of 2.3×10^9^ vector genomes at a total volume of 1µl per deposit site. All Infusions were performed at a rate of 0.2ml/min, and the needle was left in place for an additional 3-min period before it was slowly retracted. Post-surgery, the wound was closed using surgical staples and the animal was placed on a heating mat until awake.

### Stereotaxic injection of hESC transplants

For the transplantation experiments, nude rats received a total dose of 150 000 hESCs-derived neurons into striatum in a volume of 2 μl, at a concentration of 75 000 cells/μl, to the following coordinates relative to bregma: A/P: 0.5; M/L: -3; D/V (from dura) -4.5; adjusted to flat head.

### Tissue processing

Eight weeks post injection, brains were processed according to subsequent *post mortem* analysis. For RNA extractions, animals were sacrificed using CO_2_ the brains were removed and sliced in the coronal axis into two-millimeter-thick slices using a brain mold. The striatal tissue, orbitofrontal cortex, thalamus and midbrain region were rapidly dissected and frozen individually on dry ice and stored at -80 °C until RNA extraction. For immunohistochemical analysis, the animals were deeply anesthetized by sodium pentobarbital overdose (Apoteksbolaget, Sweden) and transcardially perfused with 50ml physiological saline solution followed by 250ml of freshly prepared, ice-cold, 4% paraformaldehyde (PFA) in 0.1M phosphate buffer adjusted to pH 7.4. The brains were then removed and post-fixed further for 2h in cold PFA before storing in 25% buffered sucrose for cryoprotection over at least 24h until further processing. The remaining PFA fixed brains were cut into 35mm thick coronal sections using a freezing microtome (Leica SM2000R) and collected into 8 series and stored in anti-freeze solution (0.5M sodium phosphate buffer, 30% glycerol and 30% ethylene glycol) at -20°C.

### Immunohistochemistry

For immunohistochemical analysis, tissue sections were washed (3x) with TBS (pH 7.4) and incubated for one hour in 3% H<sub>2</sub>O<sub>2</sub> in 0,5% TBS Triton solution in order to quench endogenous peroxidase activity and to increase tissue permeability. Following another washing step, the sections were blocked in 5% bovine serum and incubated for one hour and subsequently incubated with primary monoclonal antibodies overnight. To evaluate GFP expression, immunohistochemistry was performed on brain sections using chicken anti-GFP primary antibody (1:20000; ab13970, Abcam), rM3Ds expressing neurons were identified by staining for the HA-tag (mouse anti-HA, Covance Research Products Inc Cat# MMS-101R-200 RRID:AB_10064220, 1:2000). hM3Ds expressing neurons were identified by staining for mCherry (goat anti-mCherry, LifeSpan Biosciences Cat#LS-C204207, 1:1000) Following overnight incubation, the primary antibody was washed away using TBS (x3) and then incubated with secondary antibodies for two hours. For 3, 30-di-aminobenzidine (DAB) immunohistochemistry, biotinylated anti-mouse (Vector Laboratories Cat# BA-2001 RRID:AB_2336180, 1:250), anti-goat (Jackson ImmunoResearch Labs Cat# 705-065-147 RRID:AB_2340397, 1:250) and anti-chicken (Vector Laboratories Cat# BA-9010 RRID:AB_2336114, 1:250) secondary antibodies were used. The ABC-kit (Vectorlabs) was used following incubation of the secondary antibody to amplify the staining intensity through streptavidin-peroxidase conjugation and followed by color exposure in 0.01% H_2_O_2_ For fluorescence microscopy analysis, biotinylated goat anti-chicken BA9010, Vector laboratories) was used.

### Immunocytochemistry of primary glial and terminally differentiated neurons

Analysis of vector transduction efficiency in primary glial and neurons differentiated from hESC utilized immunofluorescence detection. First the medium was removed, and the cells were washed in 1x PBS. Then 100µl of 4% PFA was added to each well containing the cells and incubated at 37°C for 10 min. Following incubation, the cells were washed with PBS. The fixed cell culture was then blocked for one hour in room temperature using 100µl blocking solution consisting of KPBS with 5% BSA and 0.25 % triton-X per well. The blocking solution was the removed and replaced with primary antibody in PBS and incubated overnight at 4 degrees. Glial cells were identified using the following antibodies: rabbit anti-GFAP (1:1000; ab7260, Abcam) and Rabbit anti-IBA-1 (1:2000; 019-19741, Wako) Following overnight incubation the wells were washed twice with KPBS and then incubated with secondary antibodies in KPBS for a total of two hours in room temperature. Secondary antibodies used included: Alexa conjugated anti-rabbit (Jackson ImmunoResearch Labs Cat# 711-165-152 RRID:AB_2307443, 1:250). Finally, the cells were washed twice in KPBS and left in KBPS solution for image analysis.

### Laser scanning confocal microscopy

All immunofluorescence analysis was performed using the Leica SP8 microscope. Confocal images were always captured using a HyD detector with the lasers set to be activated in sequential mode, thus avoiding fluorescence signal bleed through. Solid-state lasers at wavelengths of 405, 488, 552 and 650nm were utilized to excite the respective fluorophores. The pinhole was always retained at Airy 1 for all image acquisitions. Post-acquisition, deconvolution was performed using the “Deconvolution” plugin for ImageJ (developed by the Biomedical Imaging Group [BIG] - EPFL – Switzerland http://bigwww.epfl.ch/) utilizing the Richardson-Lucy algorithm and applying point-spreads functions (PSFs) calculated for the specific imaging equipment using the Gibson and Lanni model in the PSF Generator (BIG, EPFL – Switzerland http://bigwww.epfl.ch/algorithms/psfgenerator/).

### Conditioned place preference test

In order to assess selective DREADD activation of the BLA on conditioned place aversion, a two-chamber box was used where each chamber was separated by a wall with a closed door. Each chamber was made different from the other using visual ques on the walls and tactile sensation on the chamber floors while retaining the same light intensity. All tests were recorded using infrared illuminated CCD cameras and the animal’s position recorded using the Stoelting ANY-maze 5.2 software package. On day one the animals were first acclimatized to one of the two chambers for a total of three hours following saline injection, being alternated within the group between the two chambers to control for any chamber bias. This trial is denoted “Control”. On day two the animals were placed in the opposite chamber from day one following s.c. injection with CNO (3mg/kg) and left in the chamber for three hours. This trial is labeled the “Conditioning” trial. On day three the door separating the chambers was removed and the animal was placed inside the box without any drug administration and recorded for a total of three hours for any apparent preferred chamber conditioning, denoted the “Preference test”.

### Elevated Plus Maze

To assay anxiety levels following selected activation of the BLA using DREADDs, the Elevated Plus Maze (EPM) was used. All animals paced in the EPM was recorded using Stoelting ANY-maze 5.2 software. The EPM was made out of black Plexiglas and consisted of four arms in the shape of a cross. Two of the arms (opposite of each other were open i.e., without walls while the remaining two arms were enclosed, i.e., had walls. The animals were injected with CNO (3mg/kg) at one and a half hour prior to the start of the test. At the start of the test the animals were placed in the center of the maze and allowed to freely explore the mazes open and closed arms while being recorded for a total of 5 min. The animals time exploring either the open or closed arms were then quantified from the recordings in order to determine the animals’ anxiety level.

### Sequencing AAV plasmid library

To facilitate paired-end Illumina sequencing of the plasmid library, a part of the AAV plasmid was excised by Cre-recombinase to bring inserted peptide sequence and barcode closer together (5a in Fig. 1B). 1.5µg DNA was incubated with 6U Cre-recombinase (NEB) in a volume of 100µl at 37°C for 1h. The reaction was terminated at 70°C for 10 min and purified by DNA Clean & Concentrator-5 (ZYMO Research). The product was digested using BsiWI and MunI, ran on agarose gel and the desired fragment was selected and purified using Zymoclean Gel DNA Recovery (Zymo Research). The gel extraction product was subjected to PreCR Repair using PreCR Repair Mix (NEB). In 50µl reaction, 50ng DNA, 100µM dNTPs and 1X NAD^+^ was incubated in 1X ThermoPol Buffer at 37°C for 20 min. 5µl PreCR repaired DNA was PCR:ed with P5/P7 Illumina primers P11 and mix of P12, P13, P14,P15 (for primers see Supplementary Fig. S13) using Phusion HSII (Thermo Fischer Scientific). Again, to reduce recombination between the fragments in PCR, an emulsion PCR was performed. The emulsion was created as described above. The PCR cycles were 1 cycle of 30s at 98°C, 18 cycles of 5s at 98°C, 15s at 63°C and 3min (8-fold of regular extension time) at 72°C, and 1 cycle of 5min at 72°C. The emulsion was broken, the product was purified as described above and a PreCR Repair was performed. 5µl PreCR repaired DNA from the previous step was used in the next emulsion PCR to add Nextera XT Indexes, using Nextera XT Index Kit (Illumina). The PCR program was; 1 cycle of 1 min at 98°C, 10 cycles of 15s at 98°C, 20s at 65°C and 3min (8-fold of regular extension time) at 72°C, and 1 cycle of 5 min at 72°C. The product from the Nextera XT Index emulsion PCR was purified and size selected using SPRIselect Kit (Beckman Culter). The purified and indexed PCR products were sequenced using Illumina NextSeq Reagent Kit v2 (Illumina) with 150bp paired-end sequencing.

### Sequencing RNA-derived barcode

Total RNA was isolated from brain tissue, primary neurons and HEK293T cells using PureLink RNA Mini Kit (Thermo Fischer Scientific) according to the manufacture’s protocol. RNA samples were incubated with DNase I (NEB) to remove DNA contamination. 5µg RNA was incubated with 1 unit DNase I in 1X DNase I Reaction Buffer to a final volume of 50µl and incubated at 37°C for 10 min. Subsequently, 0.5µl of 0.5M EDTA was added and then heat inactivated at 75°C for 10 min. DNase I-treated RNA was reverse transcribed to cDNA using qScript cDNA Synthesis Kit (Quanta) according to manufacturer’s recommendations. 2µl cDNA was then amplified by PCR using primers P16 and P17 (For primer see Supplementary Fig. S13). The PCR program was 1 cycle of 30s at 98°C, 35 cycles of 5s at 98°C, 15s at 65°C and 30s at 72°C followed by 1 cycle of 5 min at 72°C. The PCR products containing the barcodes were purified by gel extraction using Zymoclean Gel DNA Recovery Kit (Zymo Research). 20ng purified DNA was subjected to a P5/P7 Illumina adapter PCR using primers P18 and an equal mix of P12, P13, P14, P15. The PCR cycles were 1 cycle of 30s at 98°C, 10 cycles of 5s at 98°C, 15s at 65°C and 30s at 72°C, followed by 1 cycle of 5 min at 72°C. The PCR products were purified by gel extraction as previously described. Subsequently, a Nextera XT Index PCR using Nextera XT Index Kit (Illumina) was performed. The PCR program was; 1 cycle of 1 min at 98°C, 6 cycles of 15s at 98°C, 20s at 65°C and 1 min at 72°C, followed by 1 cycle of 5 min at 72°C. The PCR products were purified using SPRIselect (Beckman Culter). The purified PCR products were sequenced using Illumina NextSeq 500/550 Mid Output Kit v2 (Illumina) with 75bp paired end reads.

### Sequencing viral library

Two additional AAV batches were produced (for production method see “AAV production-capsid validation studies”), one batch with 100-fold dilution (30cpc) of plasmid containing the capsids and barcode and one 1000-fold dilution (3cpc) of the same plasmid (corresponding to 250ng and 25ng in “AAV production-Library”). After production, purification and titration, both batches were DNase I treated and lyzed by Proteinase K. The viral lysate was subjected to two rounds of PCR to add Illumina compatible P5/P7 sequences and NexteraXT Indexes and then purified using SPRIselect (Beckman Culter). The purified and indexed samples were sequenced using Illumina NextSeq500/550 Mid Output Kit v2 (Illumina) with 75bp paired end reads.

### Transduction of human ES cells & human primary Glia

Human ES cells were differentiated into dopaminergic progenitor cells^22^. 42 days after the differentiation started, cells were transduced with scAAV-GFP using 5×10^8^gc/well and incubated o.n. 72h post transduction, cells were fixed with 4% PFA and stained for Map-2, Tyrosine Hydroxylase and DAPI (see Immunohistochemistry). In total 29 different AAV-capsids were validated. Cells were analyzed in Cellomics, Trophos Plate runner and Confocal microscope. Human primary Glial cells was generated as previously described^43^. Cells were transduced with scAAV-GFP and scAAV-mCherry using 4-7×10^10^gc/well and analyzed using confocal microscopy 48h post transduction.

### Data assessment workflow

A complete interaction free workflow was implemented using the R statistical package together with a number of packages from the Bioconductor repository. From these scripts, a number of broad–utility external applications (bbmap, Blast, Starcode^44^, bowtie2,samtools, Weblogo 3 and Hammock^11^) were called and output returned to R for further analysis. This is publicly available as a Git repository at https://bitbucket.org/MNM-LU/aav-library and as a self-sustained Docker image Bjorklund/aavlib:v0.2.

In brief; barcode and sequence identification, trimming and quality filtration was conducted using the bbmap software package^45^, which allows for kmere matching of known backbone sequences against the reads. As a vast majority of barcode reads were sequenced to the length of 20 with most barcodes of a deviating length ending up being 19 bp long (Supplementary Fig. S4), for all analysis in this study, length filtration of 18≤BC≤22 was applied.

The peptide sequence fragments were similarly isolated using the bbmap software package, but this time without any application of length restrictions and then aligned to the reference peptides using blastn. The key component of the R-based analysis framework is a parallelized implementation of the MapReduce programming philosophy^46, 47^. For more details on this process please refer to^10^. In this process Bowtie2 was first utilized to align the synthesized peptide stretches to the protein reference sequences, then blastn was used to map the sequenced fragments to the peptides and finally a purpose-built R workflow was implemented so select the pure sequencing results filtering out erroneous reads generated through template switching in the PCR based sample preparation (Supplementary Fig. S4) and to identify mutations resulting from the CustomArray oligonucleotide synthesis.

From the *in vitro* and *in vivo* samples, the AAV-derived barcodes were identified by targeted sequencing and mapped back to the respective fragments and their origin within the selected proteins (Supplementary Fig. S4). Efficacy of transport was then quantified and mapped with identification of the most efficient candidates. In parallel, the barcode count together with peptide aa sequence was fed into the Hammock tool^11^ and consensus motifs were visualized using Weblogo 3.

---

**Figure S1.**
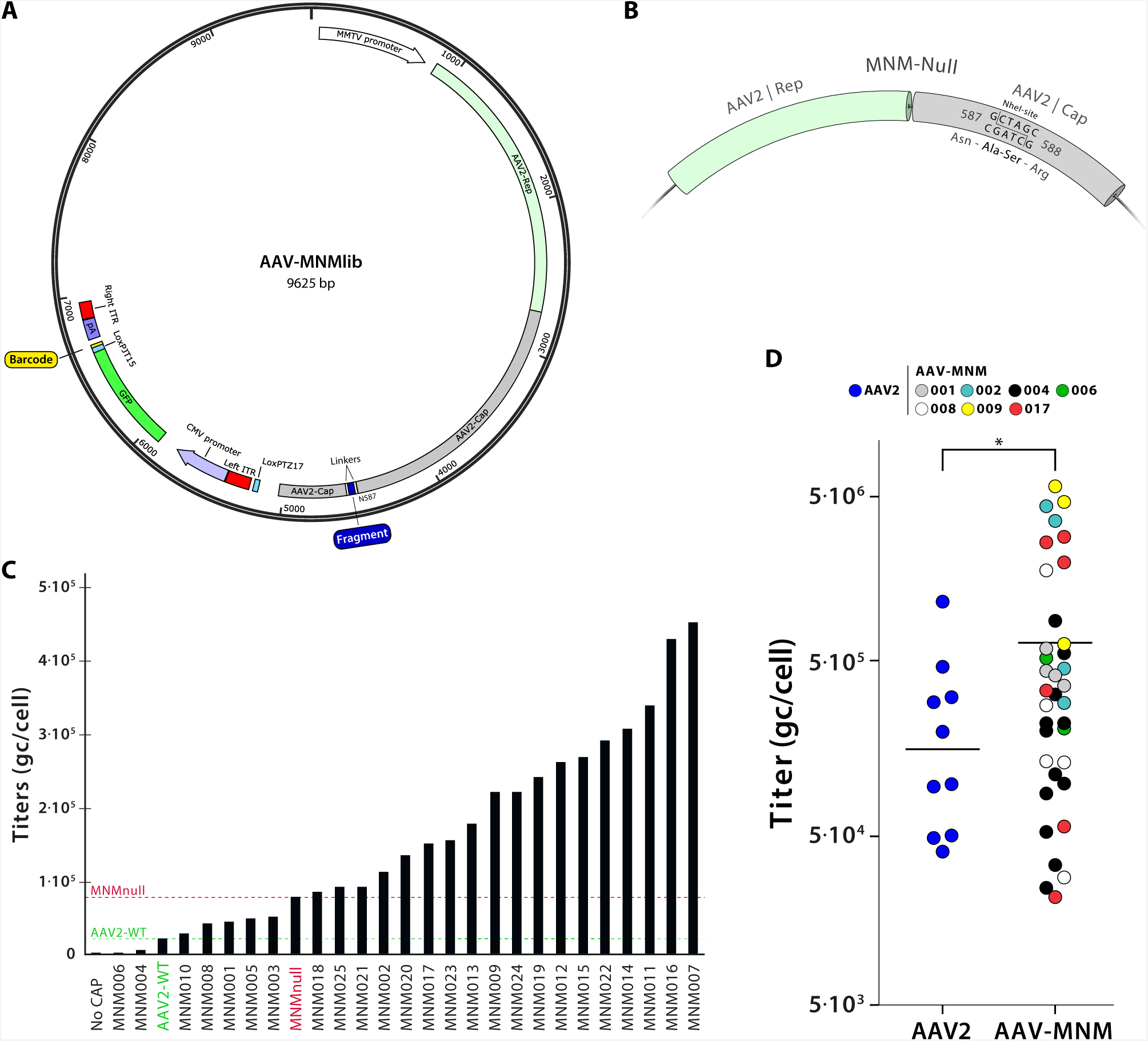
BRAVE molecular biology. **(A)** Design of the final vector for MNMlib. The mouse mammary tumor virus (MMTV) promoter is driving the expression of AAV2 Rep and Cap genes. The various fragments and linkers are inserted at the indicated N587 position of the Cap gene. Furthermore, two loxP-sites (LoxPTZ17 and LoxPJT15) are inserted to facilitate paired-end Illumina sequencing of barcodes and fragment. The barcoded expression cassette is flanked by two AAV2 inverted terminal repeats (ITRs), and thus packaged inside the formed capsids. The Cytomegalovirus (CMV) promoter is driving an expression cassette containing GFP and barcodes terminated by a SV40 poly-adenylating sequence (pA). Note that the right ITR is mutated to disrupt the nicking of the Rep protein and thus the formed genomes are self-complementary to promote rapid and efficient expression of the barcodes *in vivo*. **(B)** Design of the fragment insertion site in the AAV2 Cap gene. At amino acid position 587 a NheI site was inserted (G|CTAGC) coding for Alanine and Serine. This did not only disrupt the heparin sulfate proteoglycan receptor binding site but also facilitates Gibson assembly cloning of fragments into the Cap gene. **(C)** The graph shows titration results from a single round of virus production with AAV2-WT and selected novel serotypes (MNMnull and MNM001-MNM025, all produced in parallel with HEK293T cells of the same passage number. (**D**) The graph shows novel capsids produced at multiple occasions compared to AAV2 production. P<0.05 in two-sided T-test, line indicates the arithmetic mean. The viral titers were determined by qPCR using primers specific for the CMV promoter (see Fig. S13) and are displayed as genome copies per cell (calculated at time of transfection).

**Figure S2.**
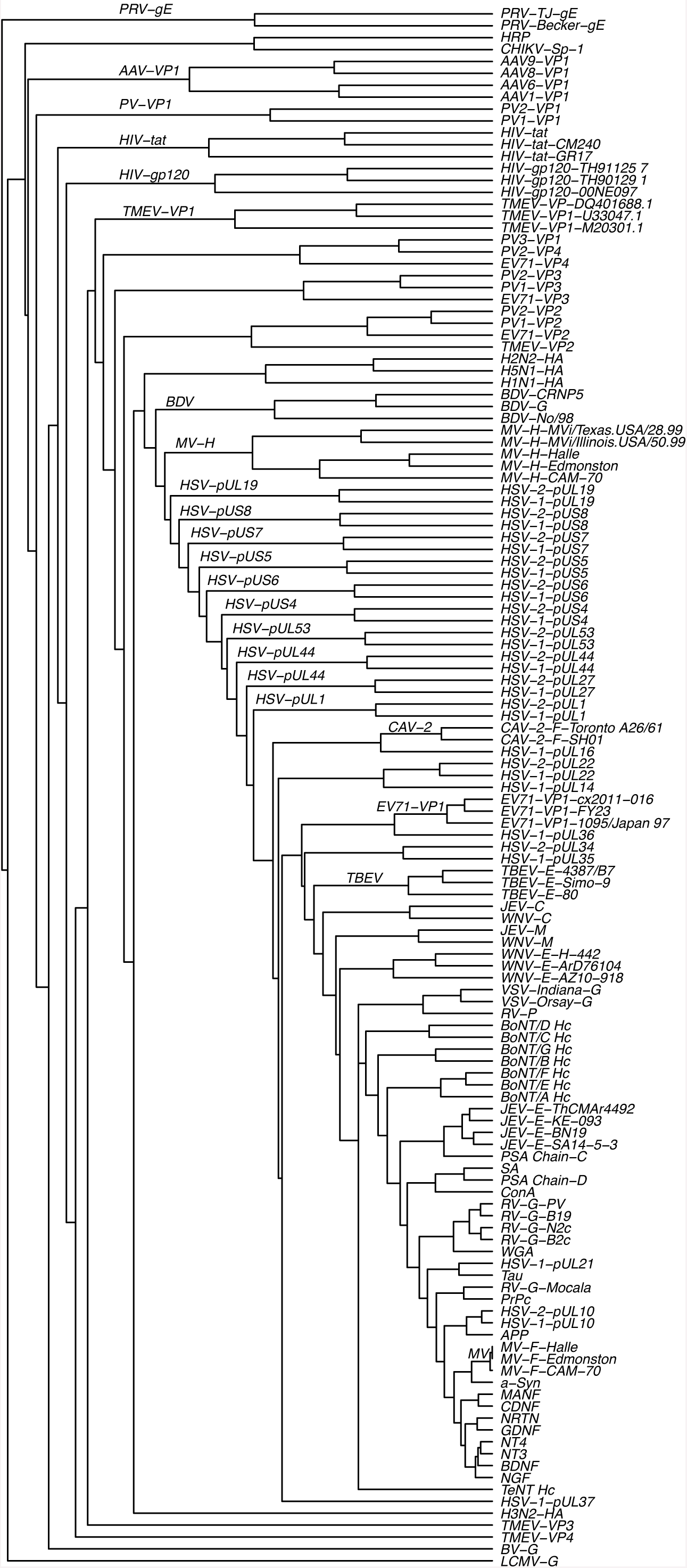
Phylotree. The amino acid sequence of all 131 proteins were fed into the USEARCH^48^ sequence analysis tool to calculate an all-pairs, hierarchical distance calculation on whole-protein similarity. The hierarchical relationship is here visualized using a dendrogram with horizontal arm length proportional to the over-all similarity.

**Figure S3.**
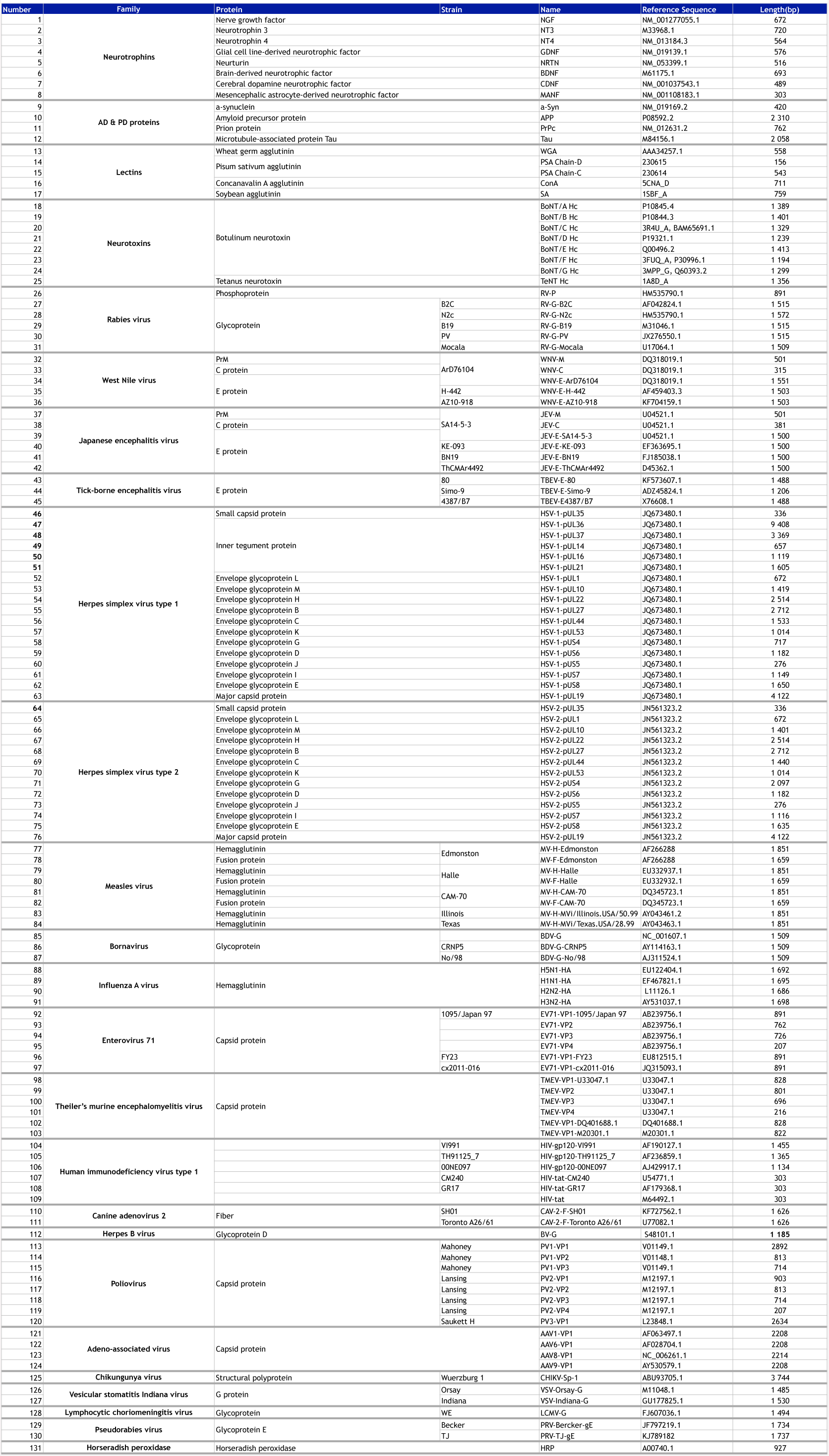
Protein Table. Summary table of all proteins included in the presented BRAVE screening experiments. All DNA sequences were downloaded from the cited reference, sequences translated into amino-acid sequences and codon optimized for expression in human cells.

**Figure S4.**
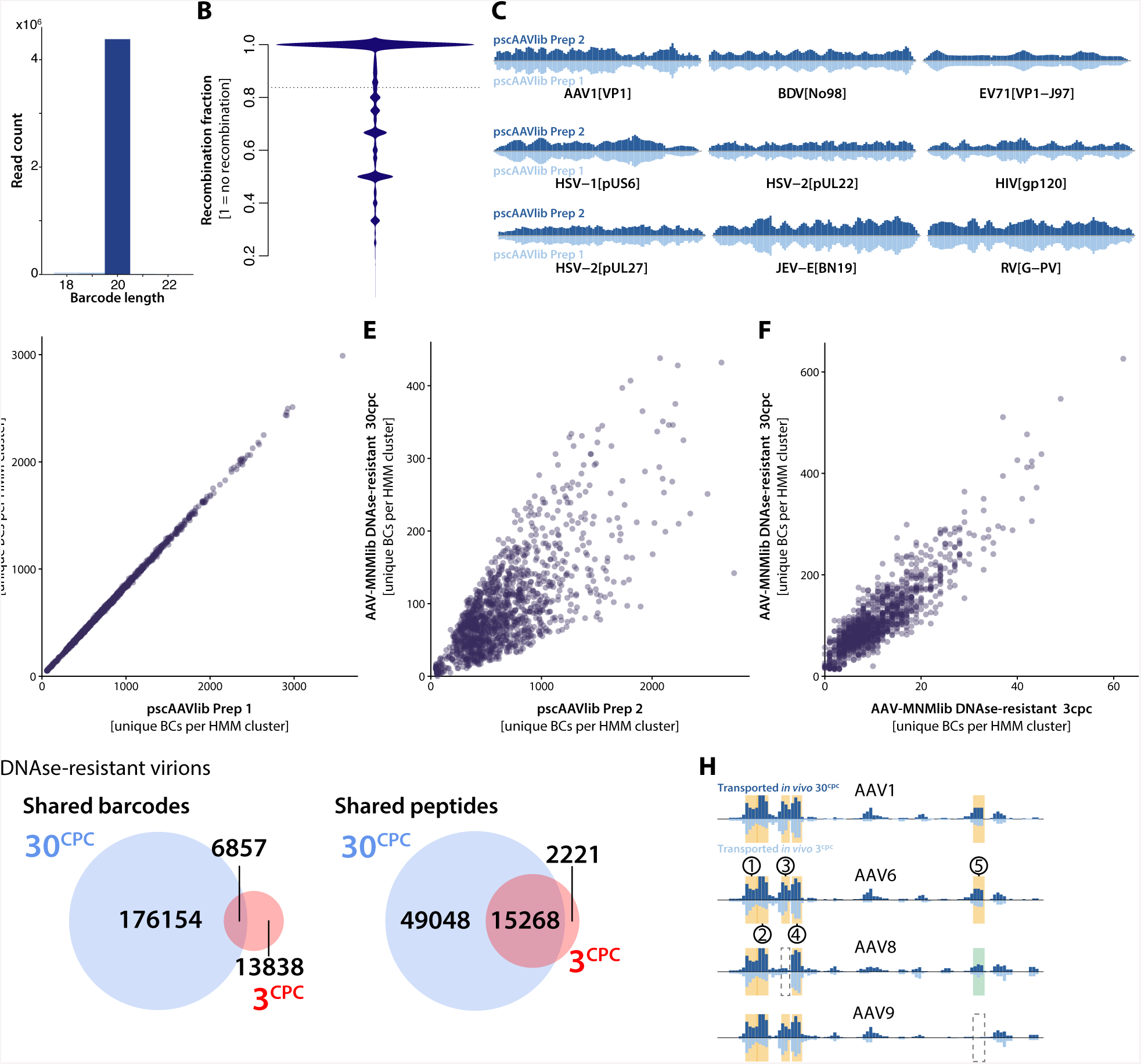
BRAVE bioinformatics. The molecular barcode was designed using a single-nucleotide exclusion cycling approach to avoid the generation of homopolymers exceeding the length of three as described previously^10^. This barcode follows the IUPAC nucleotide code: VHDBVHDBVHDBVHDBVHDB. (**A**) Through analysis of all 3.9 million unique barcodes, we could confirm that they retain the designed length and structure through production and sequencing. (**B**) To achieve cost-effective sequencing with the desired diversity, the plasmid library need to pass truncation with Cre-recombination and addition of sequencing adapters using PCR. Previously attempted approaches have resulted in significant recombination due to PCR template switching and trans-ligation between plasmids. Using emPCR and Cre-recombination we here achieve near-perfect linkage between fragment and barcode as visualized in a bean plot where 1 equals a perfect mapping between each barcode and one unique fragment. (**C**) The BRAVE approach builds on the generation of a stable bacterial library for reproducible generation of plasmid pools. Protein separated histograms confirm that the representation and diversity within all proteins is well-maintained between two plasmid pools first derived directly after transformation and expansion of the bacteria (Prep 1) and the second after a two-step secondary expansion (Prep 2). (**D**) Assessment of the relative abundance of conserved protein motifs after Hammock HMM clustering confirms a perfect preservation of the distribution between prep 1 and 2. (**E-F**) Comparing the HMM clustered motifs between the plasmid library and the DNase resistant AAV virions, we observe that most of this correlation is lost, showing that there is a function related over and underrepresentation in the packaged genomes determined by the capsid structure. Comparison of DNase resistant AAV-virions between two AAV productions rounds, however (**F**), shows an increased correlation in HMM clustered motifs, indicating that they share a majority of the functional peptides regardless of the plasmid concentration of 30cpc or 3cpc. (**G**) BC sequencing from two separate productions of AAV libraries (3cpc and 30cpc) was used to determine overlap in BCs and peptides inferred from the LUT. (**H**) A number of successful studies have utilized AAV capsid reshuffling between serotypes to generate novel capsids using directed evolution. We have here utilized BRAVE to study the potential of insertion of peptides from other AAV-serotypes into the AAV2 capsid (Fig. 3A). Interestingly the insertion of peptides from the same region of AAV1, 2 and 8 covering the N587 aa were efficiently inserted into the AAV2 capsid (Region 5 in Fig. 3A). Moreover, four additional regions (1-4 in Fig. 3A) from the N-terminal domain of VP1 (also known to be presented on the AAV capsid surface) could also be efficiently inserted.

**Figure S5.**
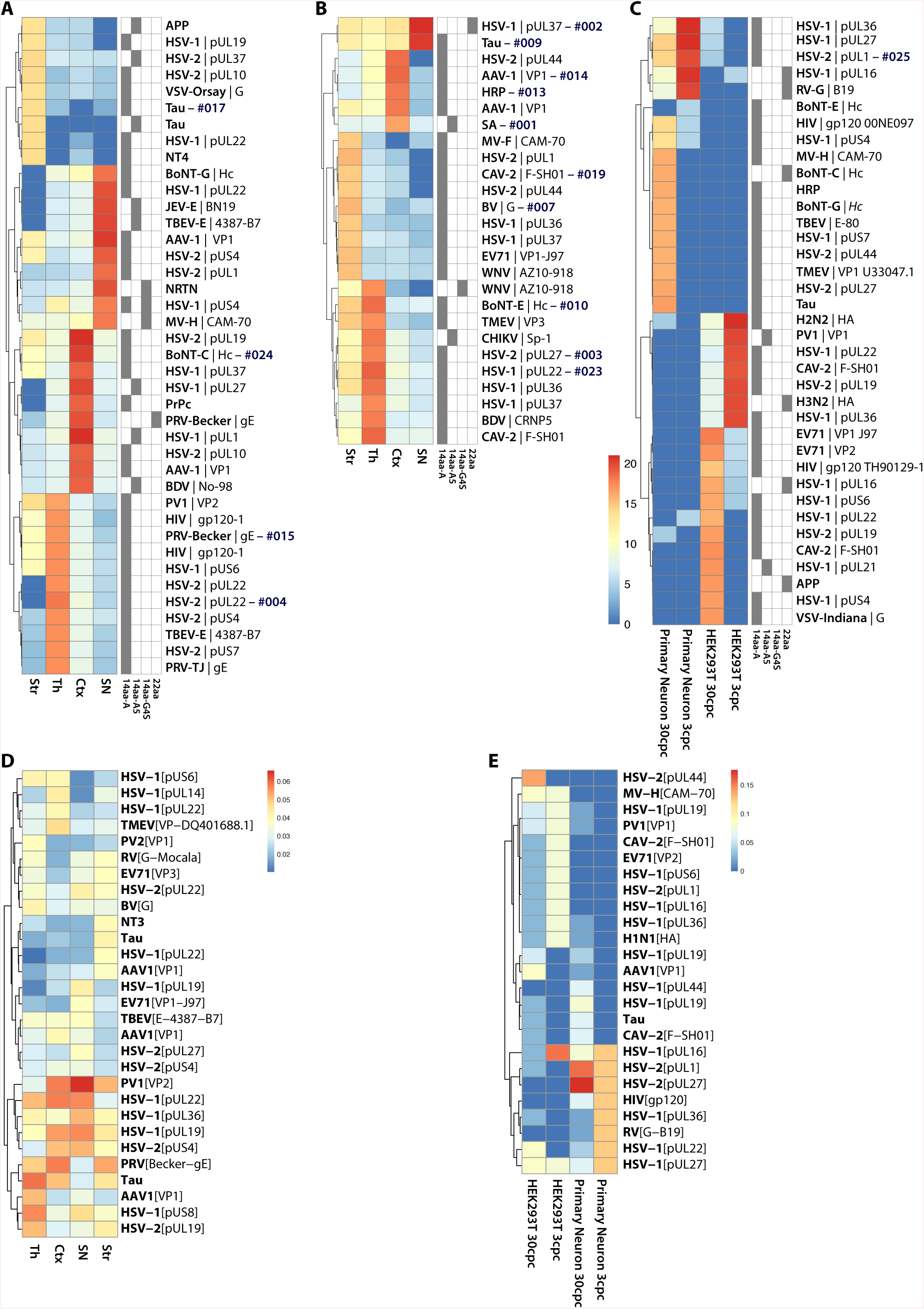
Heatmaps. To identify the most efficiently transported capsid variants, we deployed a number of alternative clustering and visualization approaches. The first is the traditional approach of relative barcode abundance in the sequencing (**A-C**). This approach provides high signal but fails to incorporate shared motifs between adjacent peptides or functional conserved motifs. Thus, we also conducted a secondary heatmap visualization using HMM clustering of the recovered peptides and incorporating a unique barcode count scoring (not including counts of the same barcode within the same region of one animal) (**D-E**). Through the selection of the top 10 sequences or motifs, we observe a higher degree of conservation when assessing the HMM clustered peptides (**D-E**) than when solely assessing each fragment individually (**A-C**).

**Figure S6.**
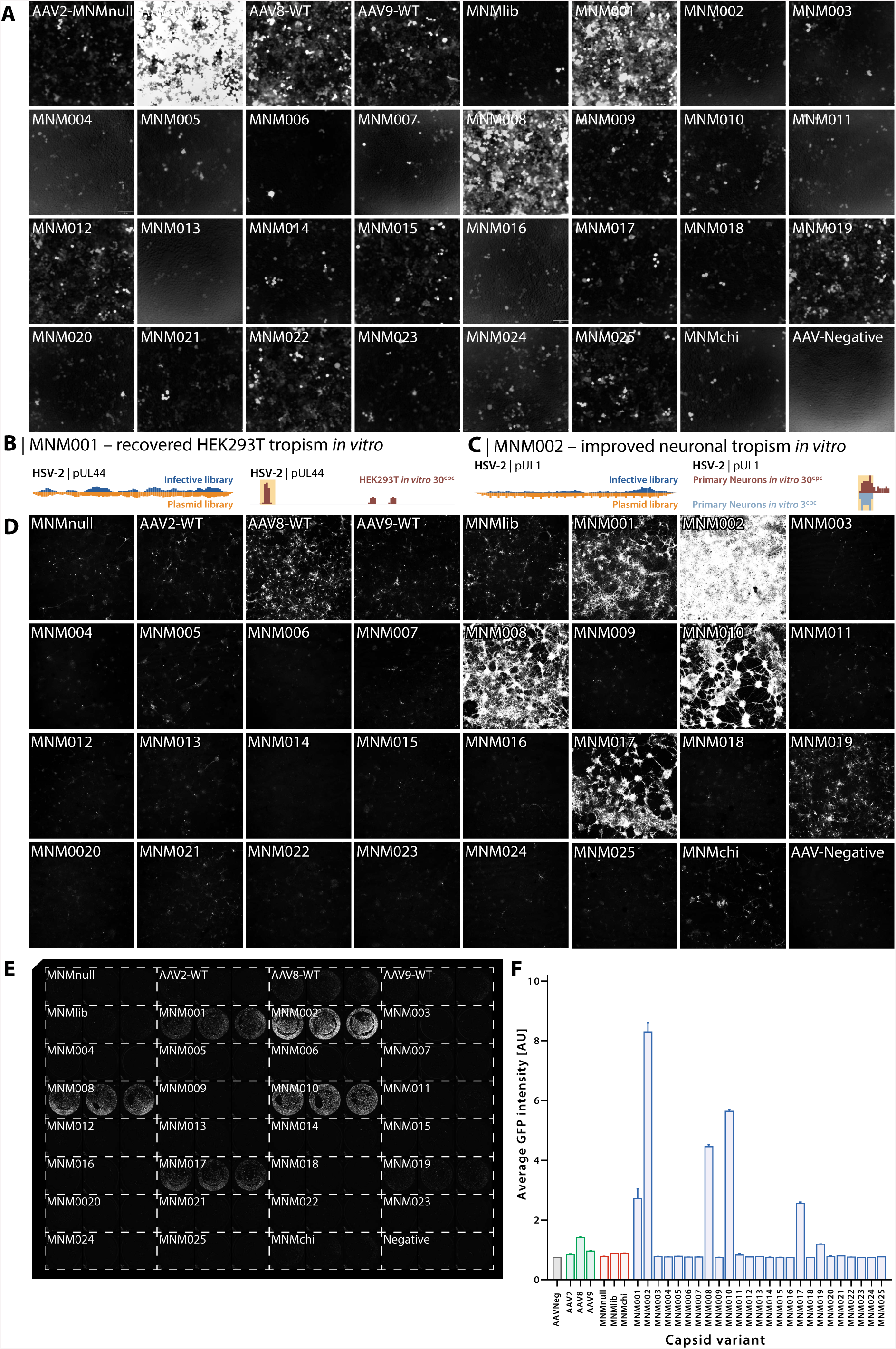
Validation of recovered HEK293T tropism and neuronal tropism in primary rat cortical neurons for BRAVE-selected capsids. **(A)** Validation of recovered HEK293T tropism for MNM001-MNM025, MNMnull, MNMlib and MNMchi compared to WTAAV2/8 & 9. All viral preps were produced to package the same scAAV-CMV-GFP genome. Cells were transduced with an MOI of 1×10^4^ and analyzed 72hours post transduction using fluorescence microscopy. Of note is that MNM001 was selected using the BRAVE approach through a screening in HEK293T cells and that MNM008 was broadly observed to have a high affinity for human cells both *in vitro* (Supplementary Fig. S11) and *in vivo* (Fig. 2). MNM001 (screened for in 293T cells) was identified from a region in the HSV-2 surface protein pUL44 (**B**) and MNM002 (screened for in primary neurons) was identified over a C-terminal region of the HSV-2 pUL1 protein (**C**). (**D**) Validation of neuronal tropism for MNM001-MNM025, MNMnull, MNMlib and MNMchi relative to AAV2/8 & 9. All viral preps were produced to package the same scAAV-CMV-GFP genome. Cells were transduced with an MOI of 1×10^4^ and analyzed 72hours post transduction using confocal microscopy and **(E)** Trophos plate runner. **(F)** Quantification of GFP expression from each triplicate in (C). The graph shows mean GFP expression and error bars represent SEM.

**Figure S7.**
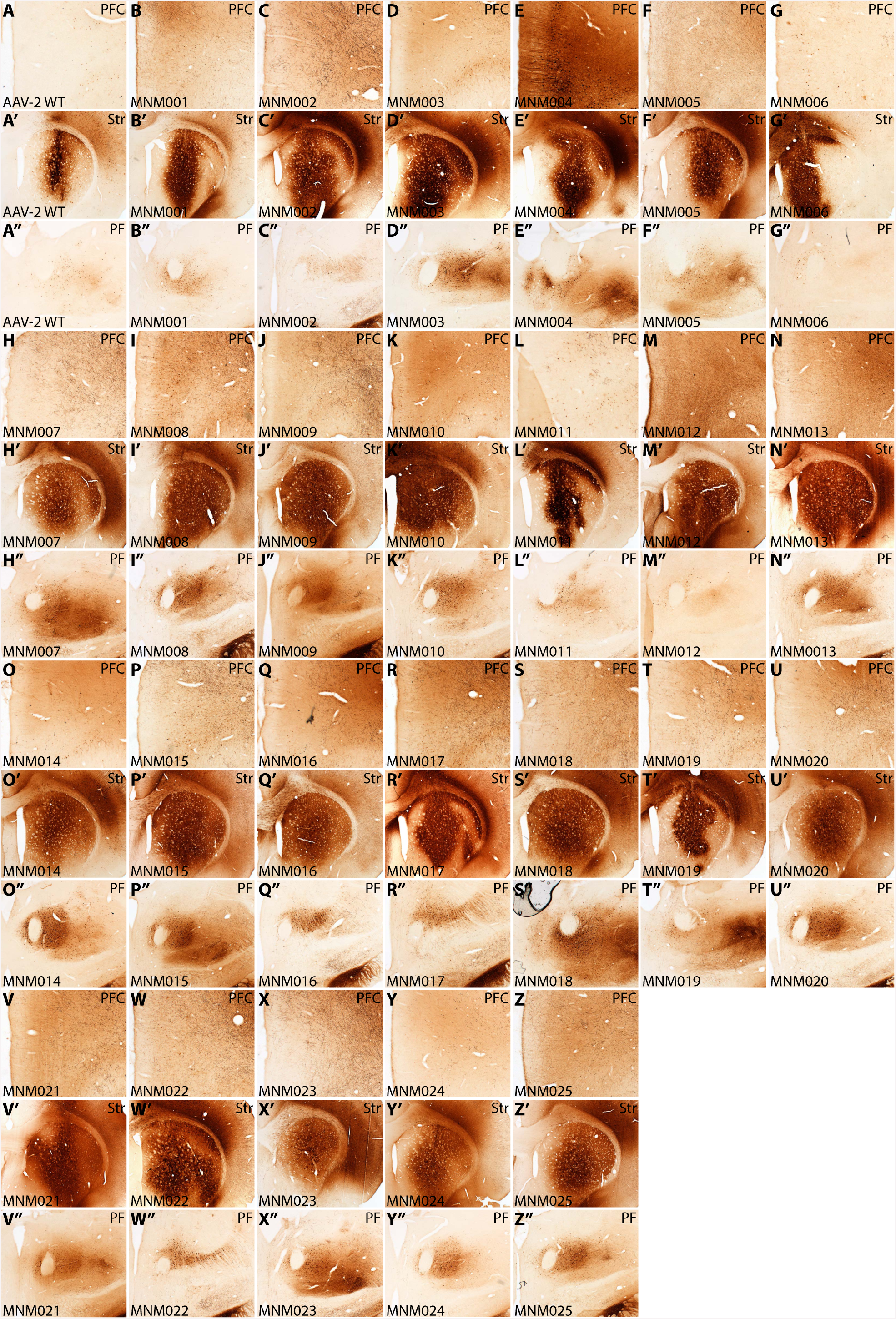
*In vivo* validation of the 25 novel capsid variants discovered using the BRAVE approach. *In vivo* validation of MNM001-MNM025 and AAV2. Vector batches produced using triple transfection with one novel capsid variant each and an scAAV genome encoding CMV-GFP was injected unilaterally in the right striatum. All vectors were diluted to 1×10^12^ gc/ml (with the exception of MNM006 which did only reach 2×10^11^ gc/ml at production, most likely due to the longer 22aa peptide inserted). Eight weeks post injection, the animals were sacrificed and the brain prepared for immunohistochemical detection of the GFP protein using a brown DAB precipitation reaction. Three 35 µm coronal brain sections were selected from each animal covering the prefontal cortex (Pfc), the parafacicular nucleus (Pf) as well as the striatum (Str), covering the injection site.

**Figure S8.**
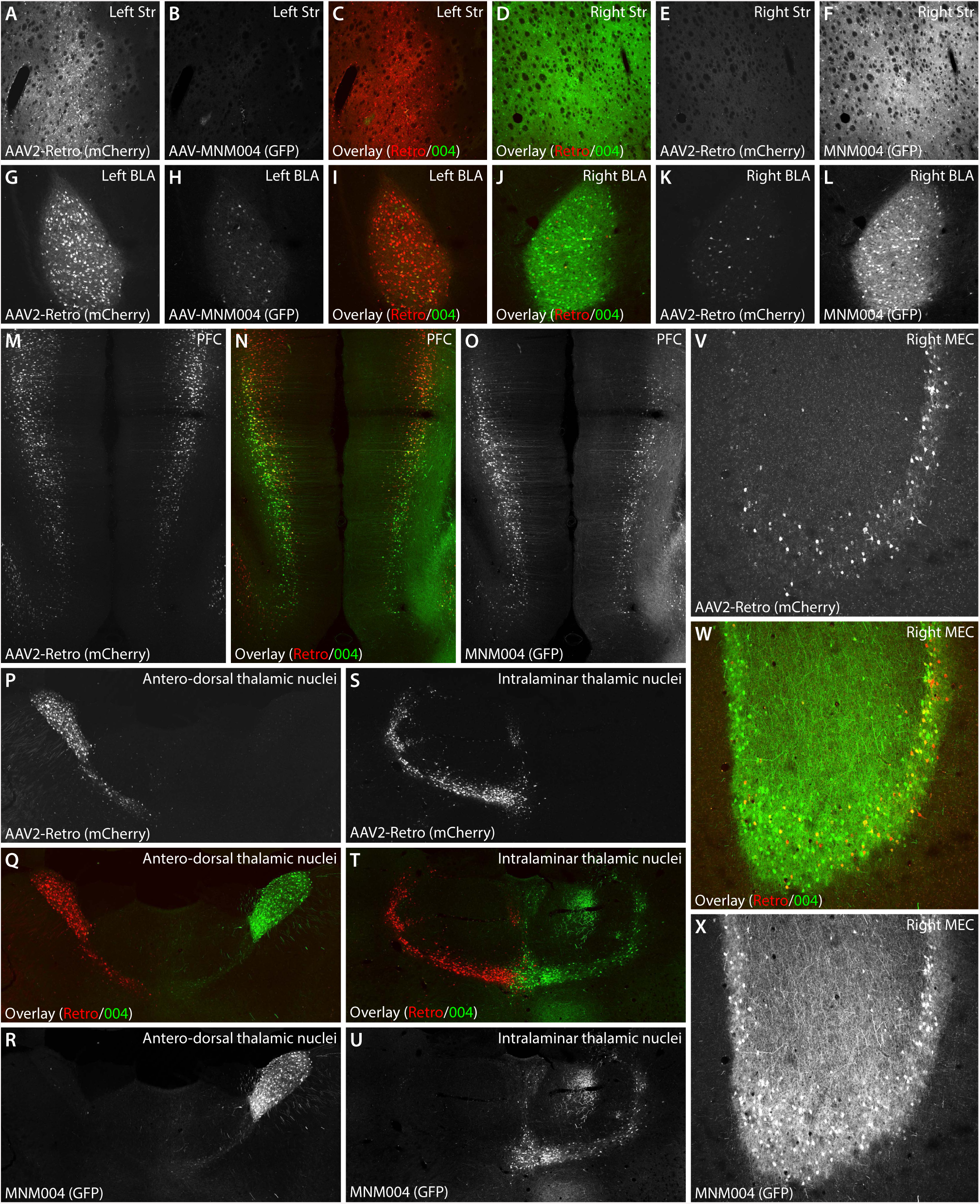
*In vivo* comparison between the MNM004 capsid variant and the AAV2-Retro peptide insertion. Validation and comparison between MNM004 and the AAV2-Retro peptide (inserted in the same backbone and position as MNM004). scAAV2-Retro[CMV-mCherry] and scMNM004[CMV-GFP] were injected on either side in the striatum in the same animal (MNM004 on the right side and AAV2-Retro on the left side) using vectors produced in parallel at the same matched titer of 1×10^12^ gc/ml. Five weeks post injection, the animals were sacrificed and brains cut into 35µm coronal sections. Selected brain areas were analyzed using laser scanning confocal microscopy visualizing the endogenous fluorescence of the GFP and mCherry to avoid antibody affinity bias and cross-reactivity.

**Figure S9.**
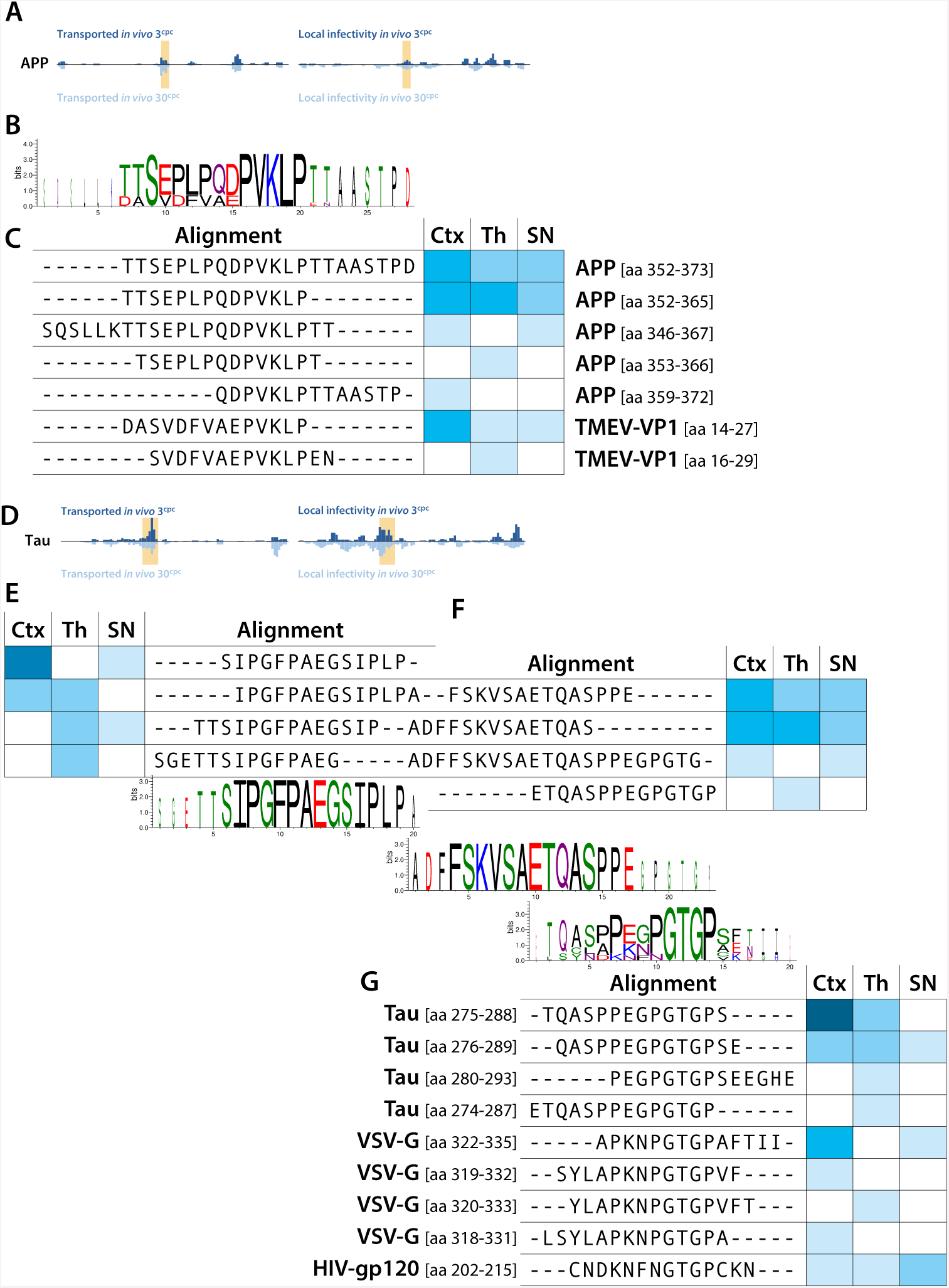
Utilization of the BRAVE approach to map proteins involved in Alzheimer’s disease *in vivo*. The BRAVE approach provides a unique possibility to systematically map protein function *in vivo*. We therefore utilized this approach to display peptides from endogenous proteins involved in Alzheimer’s disease; APP and Tau and studied if any peptides from these proteins could promote a retrograde transport of the AAV capsid and thus provide insights into the mechanism underlying the proposed cell-to-cell communication of these proteins in the disease. (**A-B**) In the mapping of APP we found two regions that conferred retrograde transports, one in the sAPP N-terminal region and one in the Amyloid beta region. (**C**) Interestingly, the sAPP region shared significant sequence homology with a region of the VP1 protein from the Theiler’s murine encephalomyelitis virus (TMEV) with appears to drive its axonal uptake and infectivity. (**D**) The functional properties of peptides originating from the microtubule associated protein Tau were even more striking. In this protein, a central region conveyed a very efficient retrograde transport. (**E-G**) This region consisted of three adjacent conserved motifs with the third motif sharing significant homology with moth the VSV-G glycoprotein (well-used to pseudotype lenti-viruses to improve neuronal tropism) and the HIV gp120 protein. Two novel capsid structures were generated from this region, AAV-MNM009 and AAV-MNM017. Both novel capsids promoted retrograde transport *in vivo* but AAV-MNM017 also displayed additional interesting properties. AAV-MNM017 infected both primary neurons and primary glial cells *in vitro* with very high efficacy.

**Figure S10.**
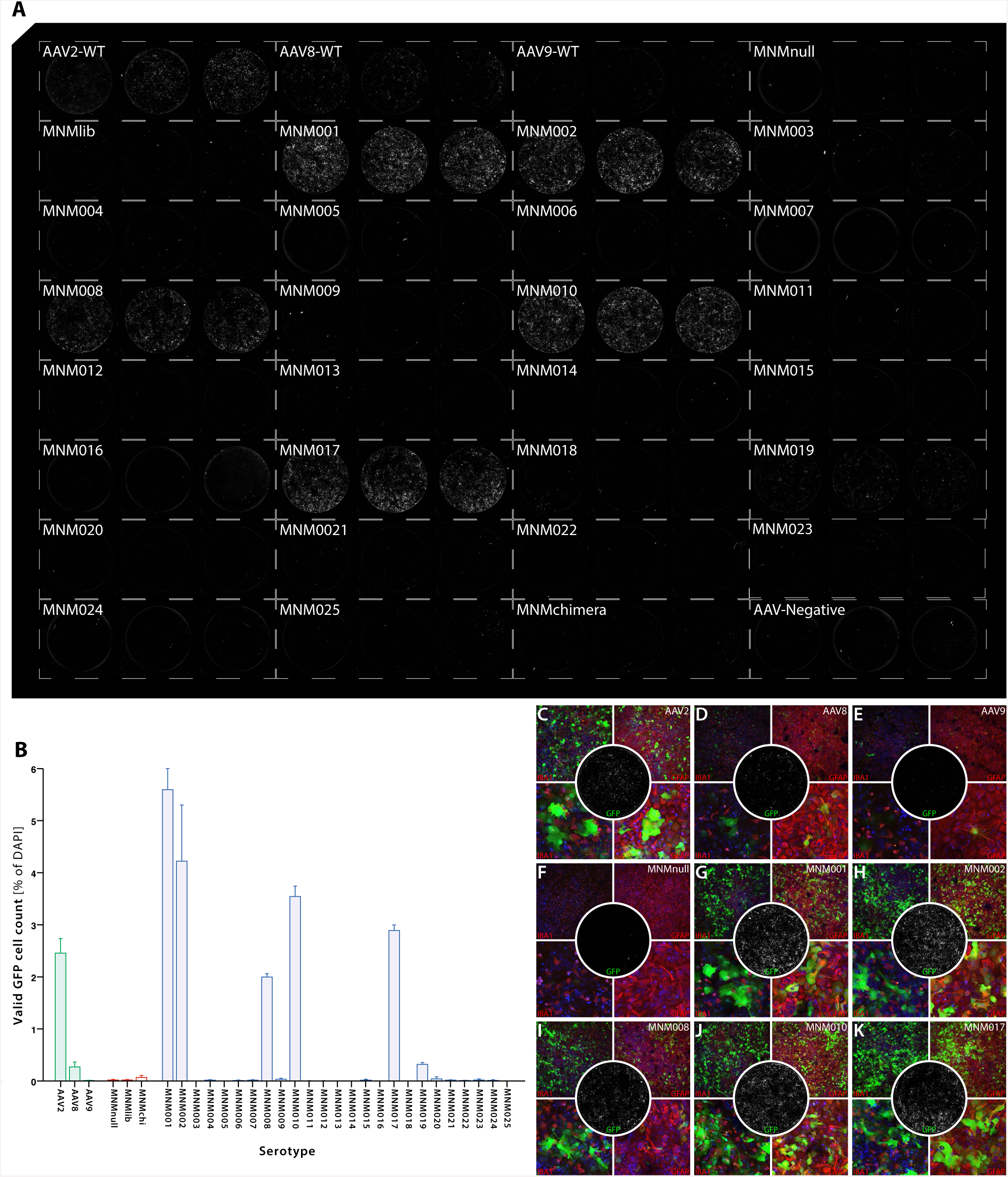
Validation of tropism in primary rat glial cells for BRAVE-selected capsids. **(A)** Validation of MNM001-MNM025, MNMnull, MNMlib and MNMChi relative to AAV2/8 & 9 in rat primary glial culture. Cells were transduced with scAAV[CMV-GFP] with an MOI of 1×10^4^and analyzed 72hours post transduction using Trophos plate runner. **(B)** Quantification of GFP expression from each triplicate using Cellomics. Graphs shows mean GFP expression and error bars represent SEM. **(C)** *In vitro* assessment of AAV2,8 and 9 and six *de novo* capsid variants in primary glial cells.

**Figure S11.**
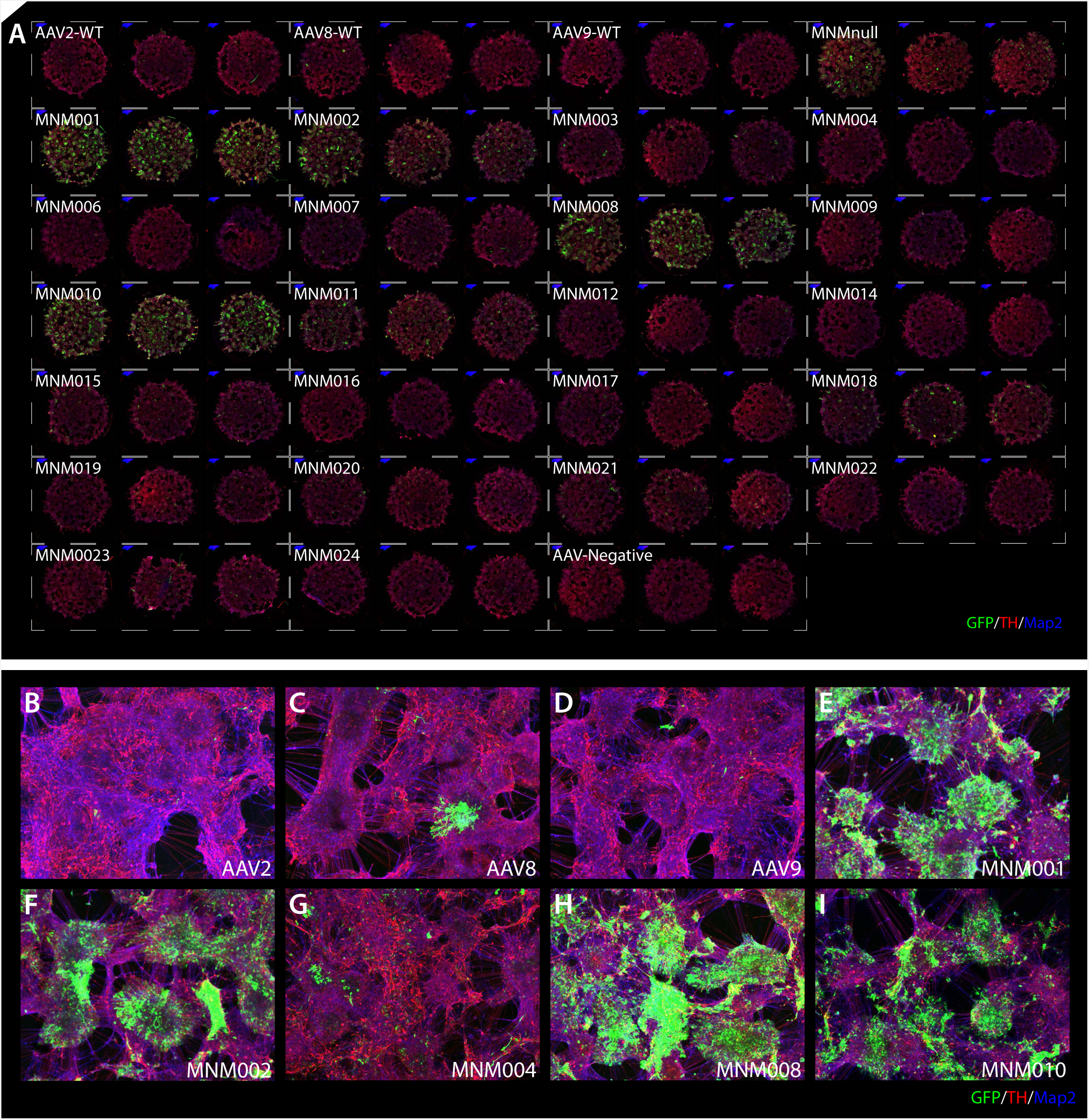
Validation of tropism in hESCs differentiated into dopamine neurons *in vitro* for BRAVE-selected capsids. (**A**) Transduction of human ES cells differentiated into dopaminergic (TH^+^) neuroblasts using selected MNM serotypes and AAV2,8&9. Cells were transduced with scAAV[CMV-GFP] with an MOI of 1×10^4^. After fixation, cells were stained for TH and Map2 (endogenous GFP expression was used for identifying infection) and analyzed using Trophos plate runner and laser scanning confocal microscopy. (**B-I**) Magnification of selected capsids from (A).

**Figure S12.**
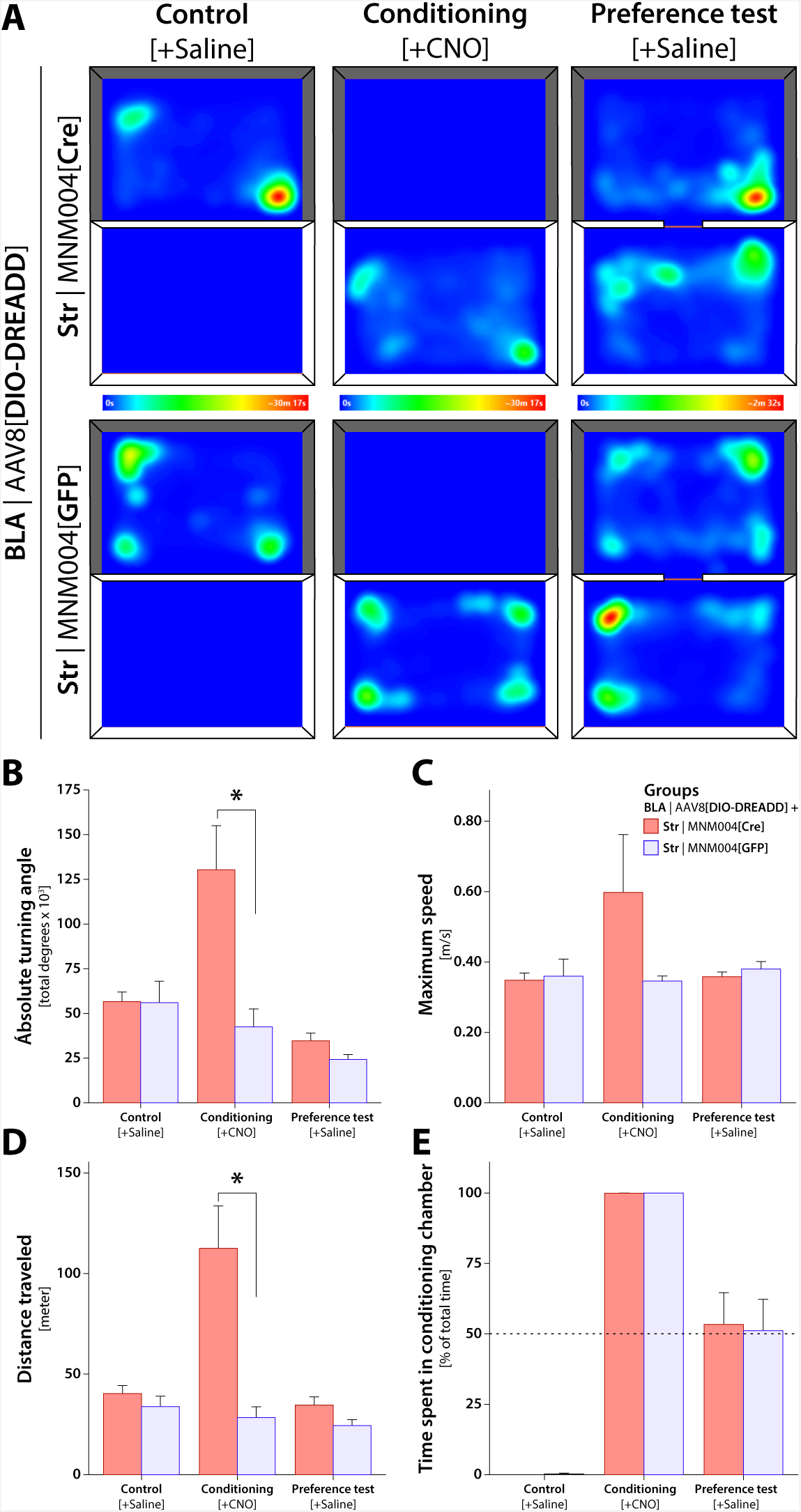
Conditioned place preference in animals with selective amygdo-striatal DREADD activation. We injected the MNM004 vector expressing Cre-recombinase in the dorsal striatum and a Cre-inducible (DIO) chemogenetic (DREADD) vector into the basolateral amygdala (BLA) bilaterally into wild-type rats. In the control animals, the Cre-recombinase in the MNM004 vector was replaced with GFP. The conditioned place preference test was used as place aversion test to study the effect of amygdo-striatal activation using chemogenetics.(**A**) The movements patterns of the two groups did not differ significantly, with all animals preferring the corners and edges over open areas. (**B-E**) After selective induction of activity of the BLA neurons projecting to the dorsal striatum using the DREADD ligand CNO, we found a striking fear and anxiety phenotype in the animals with excessive defecation, sweating a digging and freezing (see Supplementary movie 2). This was accompanied with an increase in both ipsilateral and contralateral rotation (**B**), high speed rushes (**C**) and significantly elevated mobility (**D**). However, this did not result in any conditioning, as seen in the Preference test performed the next day (**E**). Both animals in the active group and the control group spent equal time in both chamber with no signs of conditioned place aversion.

**Figure S13.**
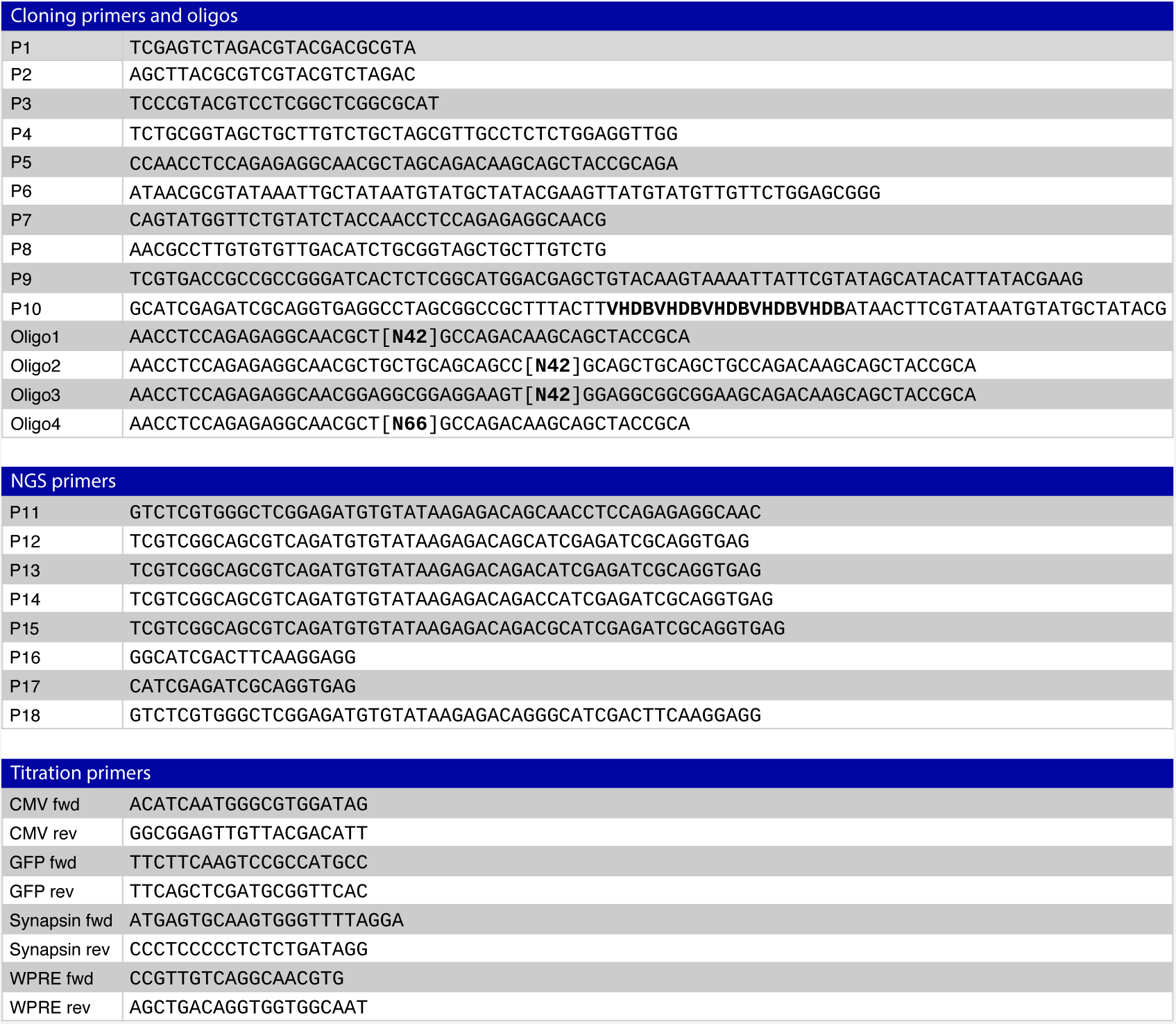
Primer table. List of most important primers used throughout the study. Primers are divided into three major groups; Cloning primers and oligos, primers for NGS and primers for viral titration. Highlighted in bold [N22] and [N66] in oligo1-4 shows the number of nucleotides to create peptides containing 14aa or 22aa. In P10, the five time repeat of VHDB (IUPAC abbreviation) highlighted in bold shows the barcode.

